# *fCite*: a fractional citation tool to quantify an individual’s scientific research output

**DOI:** 10.1101/771485

**Authors:** Lukasz Pawel Kozlowski

**Affiliations:** Solid Scientometrics Sp. z o.o., Bieliny 26-004, Holy Cross Province, Poland

**Author notes:** **ABBREVIATIONS:** RCR – Relative Citation Ratio; PMID – **P**ub**M**ed **Id**entifier; ORCID – **O**pen **R**esearcher & **C**ontributor **ID**; DORA – San Francisco **D**eclaration **o**n **R**esearch **A**ssessment; FLAE model – **f**irst-**l**ast-**a**uthor-**e**mphasis model; EC model – **e**qual **c**ontribution model; M-index – H-index divided by the number of years; fH-index – before H-index is calculated the citations are divided using FLAE model; fM-index – fH-index divided by the number of years; FLAE_RCR_ – RCR score calculated using FLAE model; EC_RCR_ – RCR score calculated using EC model; FLAE_cit_ – number of citations calculated using FLAE model; EC_cit_ – number of citations calculated using EC model; SJR – **S**CImago **J**ournal **R**ank indicator; HCR – **H**ighly **C**ited **R**esearchers list.

**Keywords:** bibliometrics, scientometrics, bibliometric analysis, quality of publications, bibliometrics tools, science impact, science policy, science evaluation, research assessment

## Abstract

Here, I present the *fCite* web service (fcite.org) a tool for the in-depth analysis of an individual’s scientific research output. While multiple existing tools (e.g., Google Scholar, iCite, Microsoft Academic) focus on the total number of citations and the H-index, I propose the analysis of the research output by considering multiple metrics to provide greater insight into a scientist’s multifaceted profile. The most distinguishing feature of *fCite* is its ability to calculate fractional scores for most of the metrics currently in use. Thanks to the division of citations (and RCR scores) by the number of authors, the tool provides a more detailed analysis of a scholar’s portfolio. *fCite* is based on PUBMED data (~18 million publications), and the statistics are calculated with respect to ORCID data (~600,000 user profiles).

## INTRODUCTION

At present, the impact of the work of a scientist can be estimated by a number of bibliometric metrics, but there is a strong bias towards the number of articles written by an author, the total number of citations of those articles, the impact factors of the journals in which they appeared (1) and finally the H-index (2). In contrast to this approach, increasing number of people are opposed to a bibliometric, mechanical *modus operandi* and in favour of expert assessment (e.g., see the DORA declaration) (3). Although expert approach is a compelling idea, in real life a fair assessment of a scientific portfolio comprising multiple publications (for instance containing over 100 items) spread across multiple journals (e.g., in 2019 PUBMED alone indexed 48,601 journals) may not be possible in a reasonable amount of time. Even if the expert is familiar with the quality of the journal in which the publications appeared and, even if he/she has read the most important publications from the portfolio, he/she still needs to understand and judge the author’s contribution to given work(s). With multiple-author papers this can be very difficult to achieve. Acknowledgment statements (if any) are usually very frugal, and it is often impossible to say which part of the work was done by which author. Moreover, if one also recognizes that publications are frequently interdisciplinary, the proper assessment of the influence of the average-sized portfolio is beyond the scope of a single person, and ultimately, it is very subjective. On the other hand, in many fields the order in the author list can be considered a rough approximation of the contribution, where the first author is the scientist who performed the most of the experiments (e.g., a PhD student), the middle authors are those who helped with multiple specialized parts of the work or/and the analysis of the data and the last/corresponding author is a principal investigator who conceived the project, obtained the funding and supervised all steps (frequently this does not exclude involvement in the experiments or the analysis). Such a model is termed the first-last-author-emphasis (FLAE) model (4), and depending on how much emphasis is placed on a particular author, the FLAE model can have multiple flavours (here we use three models named FLAE, FLEA2, and FLAE3; for details see the Material and Methods). However, given a sufficient number of items in the portfolio, they yield very similar results (5). In contrast, if the order of authors is random or alphabetical, we can always use the equal contribution (EC) model in which each author has the same weight (**Fig. 1**, Supplementary Fig. 1–4, Supplementary Tables 1–5). At this point, an open question is how to assess the influence of the publication. The most accessible (and frequently used) metric is the number of citations it has received over time. Usually, the metrics such as total citation counts or the H-index are calculated using global scores (regardless of the number of authors, each author obtains all of the citations of the publication), but applying FLAE or EC models provides a straightforward way to quantify the author’s contribution in a more precise manner. The division of the contribution is a highly demanding (and overlooked (6)) feature because the number of authors has increased steadily over time to exceed an average of six authors per research publication in 2015 (**Fig. 2**, Supplementary Table 6). The trend of having increasing numbers of authors is also clear when we analyse the mode of the number of authors in publications over the last 25 years (**Fig. 3**, Supplementary Table 7). Currently, publications with hundreds of authors are not rare, and some items can have more than several thousand authors. Concomitantly, shared first or last authorship has become a common practice, and it is not difficult to find publications with three or more shared first authors and few corresponding/last authors.

**Figure 1.**
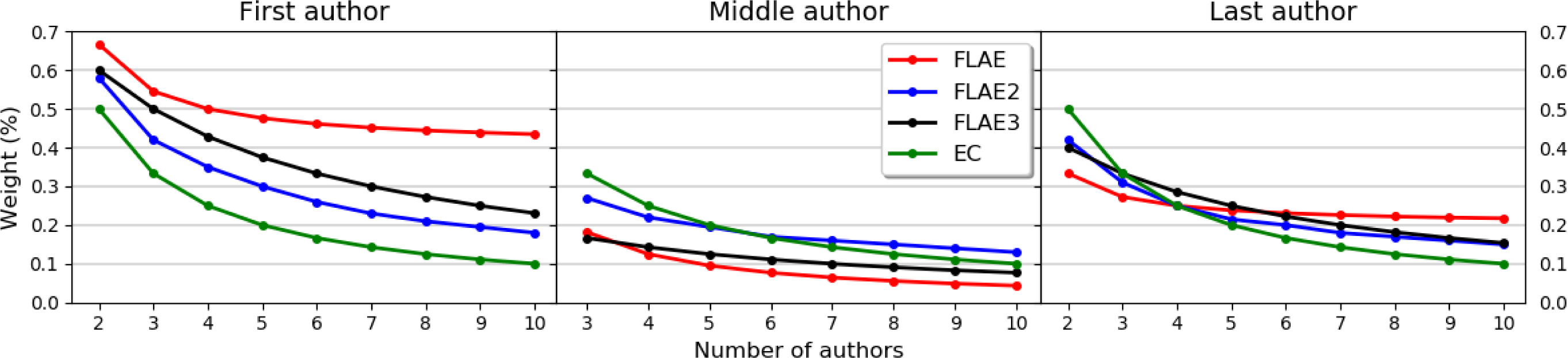
Fractional models used in *fCite* (FLAE, FLAE2, FLAE3, EC). The weights for the first, middle and last author up to ten authors. For numerical data see Supplementary Tables 1–3, respectively.

**Figure 2.**
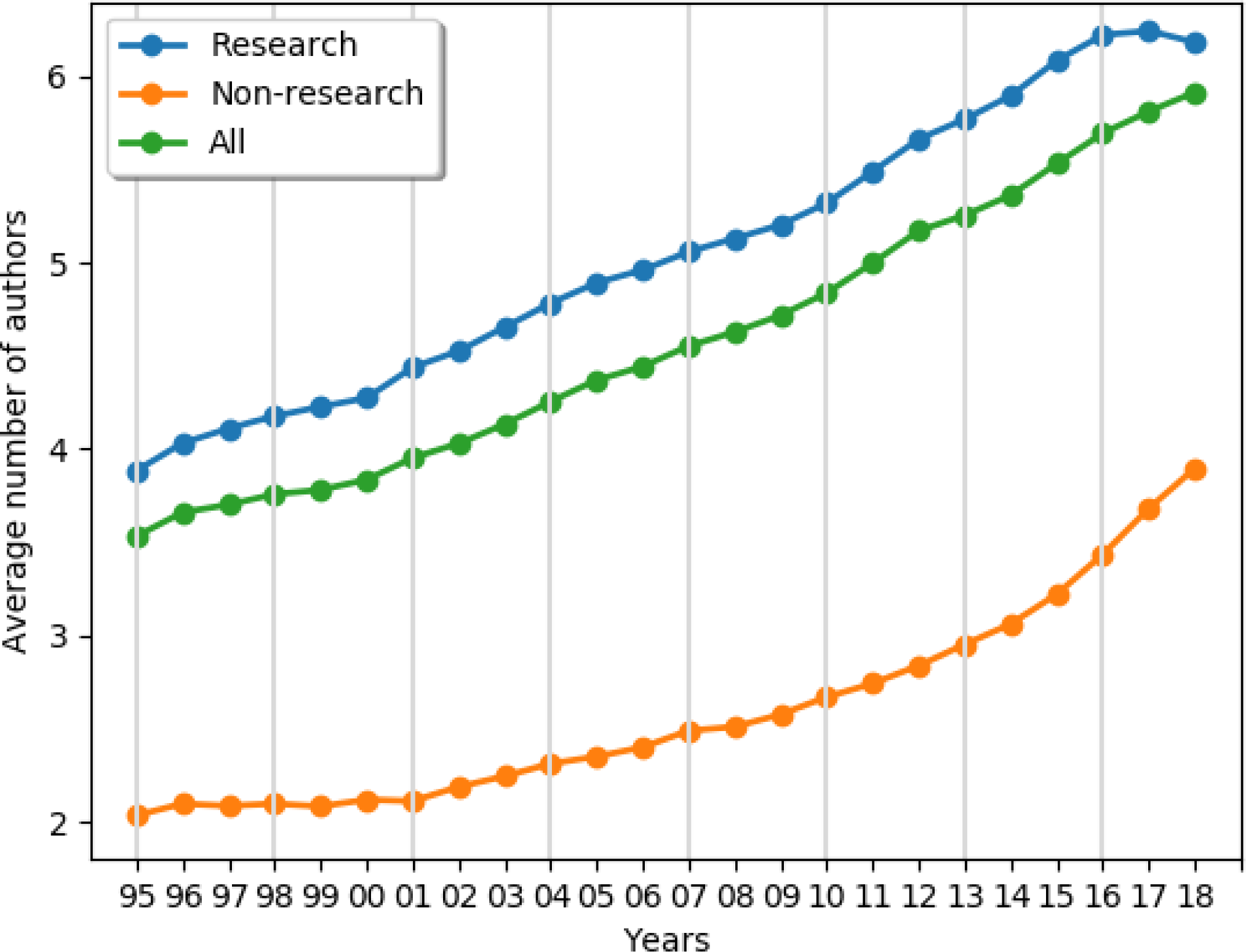
The increase in the average number of authors over time (whole PUBMED, 17 million items). For the numerical data, see Supplementary Table 6.

**Figure 3.**
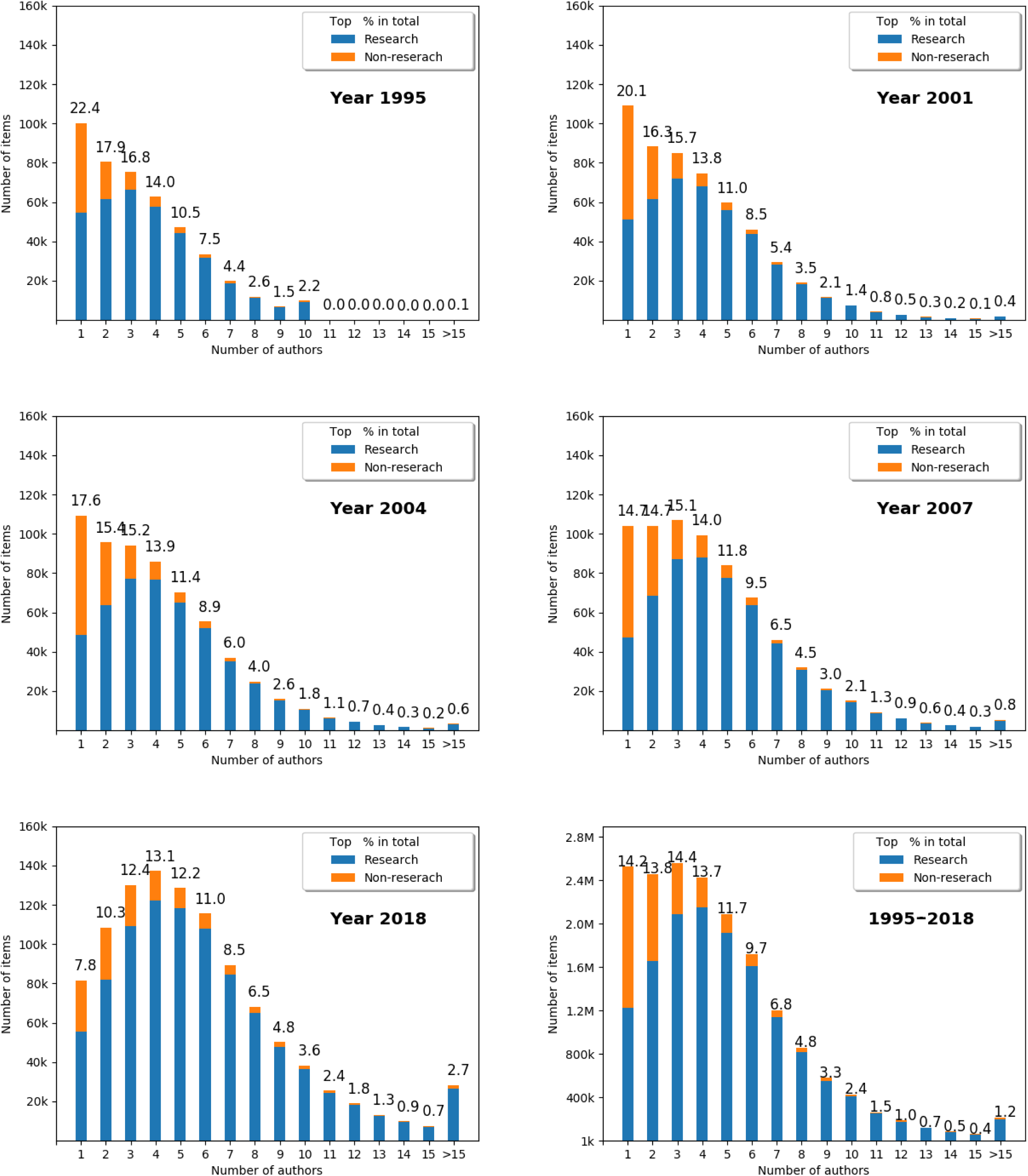
Number of authors over time with respect to research and non-research items (the years 1995-2018; 17,651,086 PUBMED publications). For the numerical data see Supplementary Table 7. Animated version of the the figure is available at: http://www.fcite.org/stats.html#Authors_over_years

Rewarding all authors, regardless of their number, is an obvious shortcoming of current bibliometric tools and contradicts Price’s law of productivity according to which the square root of the number of people is responsible for 50% of the work. This means that in 10-author publication, 3 of them do ½ of work (7). The remaining work is done by 7. Similarly, out of 100 authors, around 10 do the half of work and 90 second half. Consider a hypothetical situation in which you are a member of a grant or fellowship committee and have two applicants. Both of them published single publications in the same journal (for simplicity of the example), but the first publication has two authors (the applicant and his/her supervisor), while in the second case the applicant is the middle author of a consortium paper (such papers usually have few hundred authors). After a few years since the publication date, you see that the first publication has received a few dozen citations, while the second has a few hundred citations. Which candidate would you prefer? In the presented example, most experienced assessors would prefer the first candidate. If you were to do some back-of-the-envelope calculations, you would conclude that the first item has roughly an order of magnitude more citations per author than the second, and it is almost trivial to assess the contribution to the first publication; however, when there are a few hundred authors, it is literally impossible to say who did what, and most likely those hundreds of citations are self-citations or/and courtesy citations (for instance, the publication describes an important resource used by the whole field). The presented example is highly simplified, and usually there are more items in portfolios, which significantly complicates the analysis. Typically, portfolios will be more diverse in the number of items and journals in which they were published. The other well known drawback of the expert based evaluation system is it’s high cost (e.g, Research Excellence Framework in United Kingdom) (8). The *fCite* web service presented here should fill the gap between the (overly) simple bibliometric and expensive expert-based approaches, facilitating a fairer assessment of scientific output.

## MATERIALS AND METHODS

### Data sets

The statistics in fCite are based on two data sets: (1) PUBMED data set – contains PUBMED publications (17,787,016 publications; 14,444,982 research and 3,342,034 non-research items) obtained via icite.od.nih.gov portal (Supplementary Data 1-2) and (2) ORCID (Open Researcher & Contributor ID) data set – contains 5,380,983 user profiles (Supplementary Data 3).

### Fractional contribution models of the authorship

Four, different models had been used to assess the author contribution:

- FLAE (first-last-author-emphasis) model is based on Tscharntke et al. 2007 definition with slight modifications (4). The contribution of individual authors can be described briefly as “the first author gets 100, the last 50, and all others 100/number of authors and then scores are normalized to 1”. This type of the model gives the strongest weights to the first and the last author penalizing middle authorship (Supplementary Table 1).
- FLAE2 model is based on Corrêa Jr. et al. 2017 (5). This is empirical model based on the authorship contribution for the mega-journal PLoS ONE (~65,000 publications). On average this model is more benign for middle authors. As the data presented by Corrêa Jr. and co-authors are limited up to ten authors, for the longer author lists the contribution has been modeled by curve fitting with some noise using the initial matrix (with up to ten authors), and thresholds 0.06 for 30 authors and 0.07 for 100 authors (for more details see scipy.optimize.curve_fit documentation and http://www.fcite.org/FLAE2.txt) (Supplementary Table 2). Additionally, this model is asymmetric, i.e. middle author weights depends on the position, the closer to the first author, the better are weights for middle author (up to 10^th^ author).
- FLEA3 model is a simple variation of FLAE model, but the contribution of individual authors is more equal. It can be shortly described as “the first author always get at least three times more than co-authors, and the last author at least two times more than other co-authors” (Supplementary Table 3).
- EC (equal contribution) model assumes that each author contributed equally to given work (Supplementary Table 4).

First three models assume that for a given field the order of the authors is not random and the first author was the one who contributed the most while the last is a senior author who conceived the project (frequently the corresponding author). Such assumption is true for many sub-fields of the biomedical sciences. Alternatively, in many other sub-fields the order of the authors can be alphabetical, ordered from the most significant to the least significant author or completely random or irregular. In such cases EC model should be used. For simplicity, all models assume that there is only one first and one last author which is not necessarily true as with strong pressure for the publishing in the top journals and having more and more authors per the work, nowadays many papers have multiple first and senior authors. All four models are metrics agnostics thus they can be used for the citations, RCR scores, or/and H-index.

### Additional metrics used in *fCite*

In order to analyze the portfolio, the user is asked to provide all combination of author names and surnames, and the list of PMID (**P**ub**M**ed **ID**entifier) ids. As a result he/she obtains:

a. the size of the portfolio with the time span of the publishing period,
b. the number (and the percentage) of the single, the first, the last and the middle author papers,
c. H-index (2),
d. M-index (H-index divided by the number of years from the first publication),
e. fH-index and fM-index (the citations are divided according the author contribution to each paper using FLEA model),
f. the average number of the papers per year,
g. the total and fractional citation and RCR scores based on FLAE, FLAE2, FLAE3, EC models (Citations, total RCR, FLAE_RCR_, FLAE2_RCR_, FLAE3_RCR_, EC_RCR_, FLAE_cit_, FLAE2_cit_, FLAE3_cit_, EC_cit_, respectively),
h. the average number of the authors,
i. FLEA per year (RCR),
j. the average FLAE article score (RCR),
k. the average article impact per year (RCR),
l. the ratio between FLAE_RCR_ and total RCR and the expected value,
m. sortable table for individual publications with PMID, year, title, authors, article-type, journal, and FLAE_RCR_, FLAE2_RCR_, FLAE3_RCR_, EC_RCR_, FLAE_cit_, FLAE2_cit_, FLAE3_cit_, Ec_cit_, Citations, total RCR scores.

### Initial data cleaning

In order to analyze the authorship patterns, the PUBMED data set (over 17 million of publications) had been mapped into ORCID portfolios (over 5 million of users). The ORCID data set provided author name and surname with the list of publications. First, an empty records (the profiles without public data) had been discarded (4,217,452 out of 5,380,983 records). Next, the portfolios with at least one publication with the DOI, PMID or PMC identifiers had been filtered. This gave 1,154,443 portfolios (with 19,516,285 non-unique articles in total). As 19,097,891 (97,85%) of items had only DOI identifier, additional step was required (namely mapping DOI to PMID identifiers). The whole PUBMED records (27,414,004 publications) had been search for DOI using summary XML files and *eutils* tool provided by the National Institutes of Health (NIH). As a result 599,468 (7,813,971 articles) of non-empty portfolios with at least one PMID had been obtained.

### Example record from ORCID (csv format)

~~~
ORCID,surname,name,list_of_PMIDS
0000-0002-2518-5940,Liebovitz,David,23550982||23646091||19468082||22034582||19267397||17219478||
19647184||28527507||17219519
~~~

### Example record from PUBMED (json format)

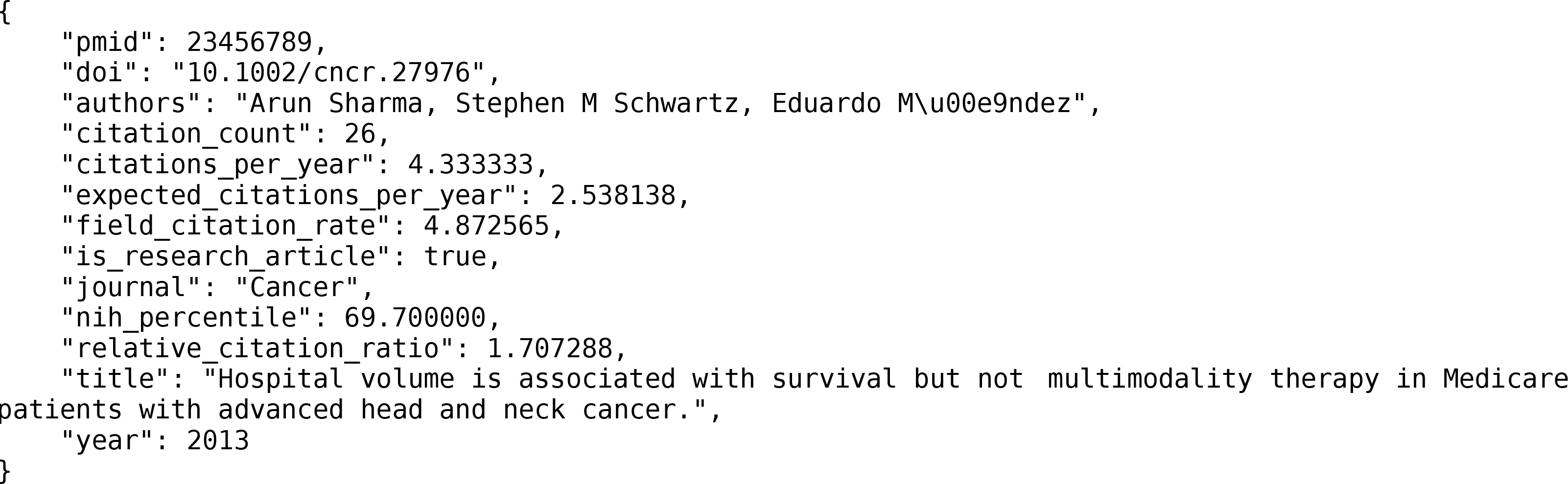

*fCite* uses following fields: authors, citation_count, relative_citation_ratio, is_research_article, year and pmid.

### Data analysis

One of the first steps of the analysis was to clean the name and surname provided by the ORCID database. The data in ORCID are in the UNICODE (UTF-8) format which means that they can contain any Non-English letters. Thus, at this step all surnames and names had been translated to equivalents of English letters (e.g., Kozłowski Łukasz to Kozlowski Lukasz, 吴锋 to Wu Feng). Then, given the list of PMIDs in the portfolio, all publication records from PUBMED had been retrieved. In order to identify the author position on the authorship list, Levenshtein and Jaro–Winkler distances had been applied in the following way. First, a set of possible surname and name combinations had been prepared (name surname, surname name, n surname, name initial surname, etc. for instance given John Smith, the set contained john smith, smith john, j smith, john x smith, etc.). This step was required as the order of the name, surname, initials and the letter size are frequently different in the databases or/and particular publication records. Next, for each author in the individual authorship list the Jaro–Winkler distance is calculated. The author which has the highest Jaro-Winkler distance is used (the similarity threshold of 0.7 is used to filter out non-important hits). From now on, for each publication in the portfolio, the position of the author is available and can be used to divide the publications into the sole, first, last and middle author ones. Having positions of the authors allow to use fractional models (FLAE, FLAE2, FLAE3 and EC) to calculate fractional scores for the citations and RCR metrics.

ORCID data had been also used to quantify the significance of obtained scores. For instance, it is not enough to say that FLAE_RCR_ (or any other score) is equal to 10. Obviously, the bigger the number the better, but it is useful to compare it to some reference. For this purpose we calculated the percentiles for the score in respect to all ORCID portfolios (the value below which a given percentage of observations in a group of observations falls).

Note that the percentiles presented by *fCite* are calculated separately for each individual metrics including division into research and non-research item portfolios (**Table 1**). Additionally, as the distribution of bibliometric metrics is practically never normal (Gaussian) (Supplementary Fig. 5), the percentiles are presented with additional precision (this allow to distinguish similar portfolios, especially top ones).

**TABLE 1.**
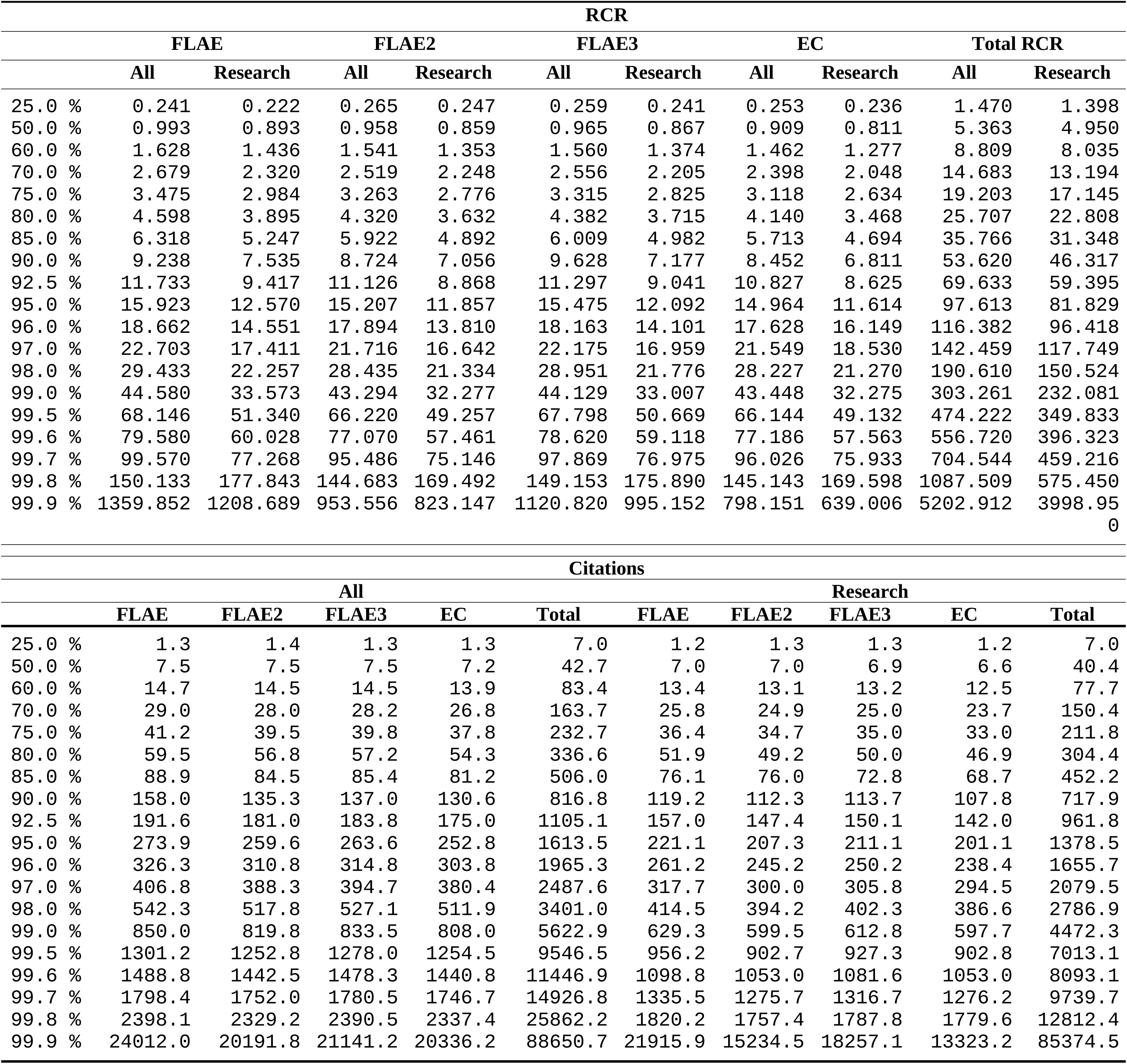
Percentile scores based on ORCID profiles in 2018 for the key metrics used in *fCite* with respect to RCR and citations (a selection of the thresholds is presented; all data were bootstrapped 1000 times; and for complete list with the supporting values, see the files at http://www.fcite.org/percentiles_2018/). parts For the ratio and spread between fractional metrics (e.g., FLEA) and total RCR (Total Citations), see Supplementary Figures 1–2.

## RESULTS

The primary result of the work presented here is, the web service (fcite.org), where an author’s contribution can be calculated using fractional models in addition to the plethora of statistics related to a given portfolio (**Fig. 4**). The *fCite* service operates using a list of PMID (**P**ub**M**ed **ID**entifier) and/or ORCID (**O**pen **R**esearcher & **C**ontributor **ID**) ids accompanied by all combinations of the names and surnames of a given author. The analysis can be performed for all items (research articles and non-research items such as editorials, reviews, and others) or separately only for the research items. The *fCite* service relies on citations and so-called RCR (**r**elative **c**itation **r**atio) scores that come from the iCite web service and are calculated based on the PUBMED database (9). As of October 2019, *fCite* comprises over 17 million publications and counting (the *fCite* database grows by ~100,000 items each month). The reference for the scores comes from the analysis of ORCID data (572,910 profiles with 7,008,012 unique publications in total). The ORCID profiles provide the names of the authors, together with the lists of co-authored publications. Thus, by using string metrics such as Levenshtein and Jaro–Winkler distances (10), it is possible to identify an author’s position in the list of the authors for each publication. This provides the unique opportunity to study authorship patterns depending on whether the person is the first, middle, last or single author. These data provide a solid foundation for the assessment of the author’s position importance with respect to portfolio size and time. As shown in **Fig. 5** (and Supplementary Tables 8–9), at the beginning of a scientist’s career (with small portfolios with fewer than 10 items), a substantial number of first author papers are expected. As the researcher progresses (and the number of publications increases), the last author publications begin to take the place of the first author publications. Surprisingly, the middle and single authored fractions are roughly stable regardless of portfolio size (where single author papers are extremely rare, and middle author papers constitute over half of the items). Moreover, calculating the main scores provided by *fCite* (e.g., FLAE_RCR_, FLAE2_RCR_, FLAE_Cit_, FLAE2_Cit_, EC_RCR_, EC_Cit,_) for the ORCID users provides solid ground for the assessment of the importance of the obtained numbers (so-called percentiles) (**Table 1**).

**Figure 4.**
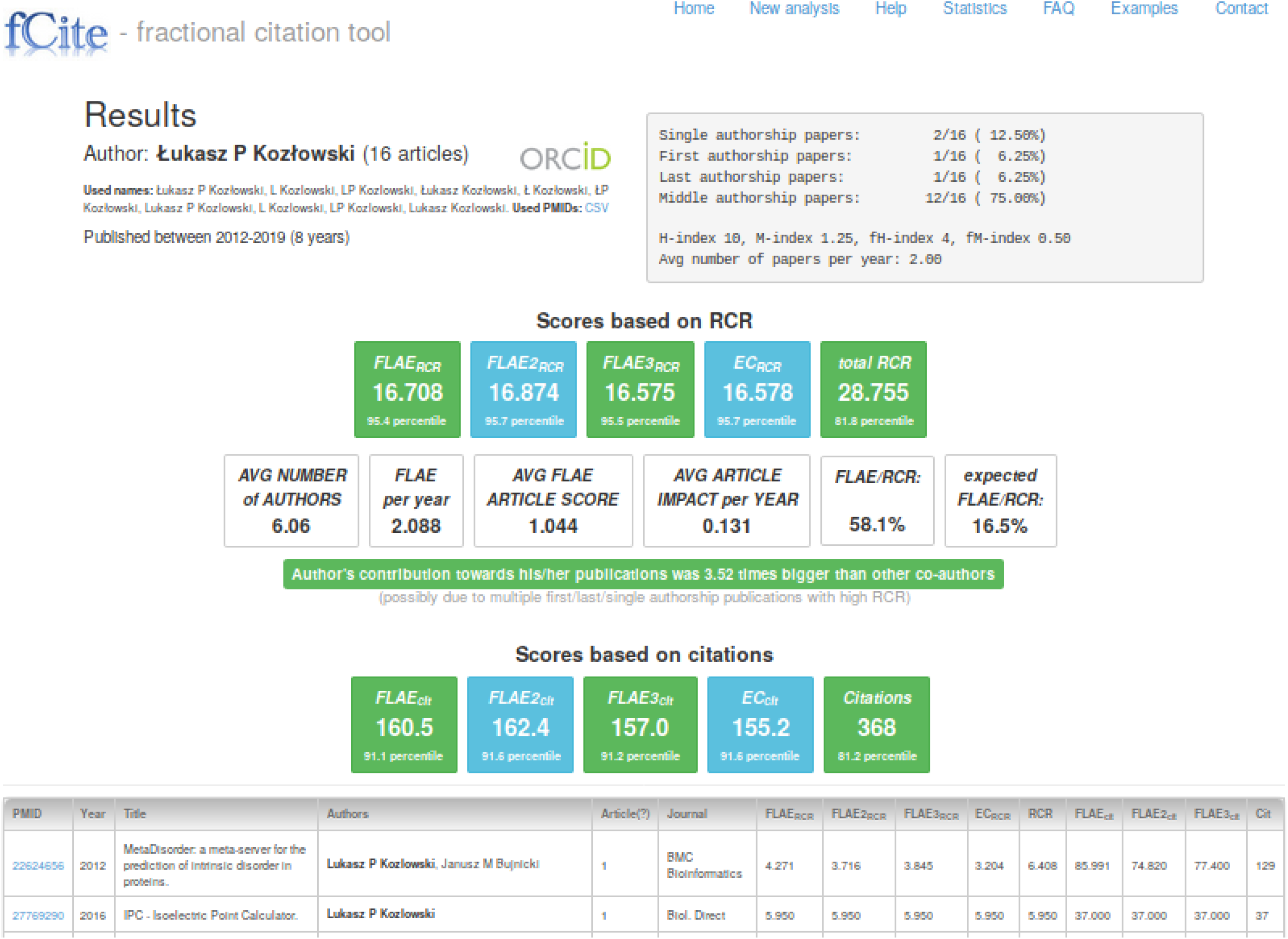
Example output from *fCite*. Given the list of PMIDs (16 items in this case) and all combinations of the name, the user obtains a detailed analysis of the researcher’s contribution (for a brief user manual for *fCite,* see the “Help” section). On the top left, you have the number of articles and the time span within they were published. Next, in the top-right panel, you have some statistics about single, first, last and middle authorship alongside the H-index, M-index and their fractional analogues. In the centre, you have the scores for the fractional models based on RCR (FLAE_RCR_, FLAE2_RCR_, FLAE3_RCR_, EC_RCR_, FLAE_RCR_ and total RCR). Then, using those values, you obtain scores such as the average number of the authors in the publication, FLAE_RCR_ per year, average article FLAE_RCR_, the article impact per year, and the ratio of FLAE_RCR_/RCR alongside the expected ratio based on the number of authors in the portfolio. The next line is a simple repetition of the scores but here based on raw citations. Note that the main scores are accompanied by the percentiles (based on ORCID portfolios) to facilitate the assessment of the importance of the scores. Finally, you have a sortable table with individual publications and their scores.

**Figure 5.**
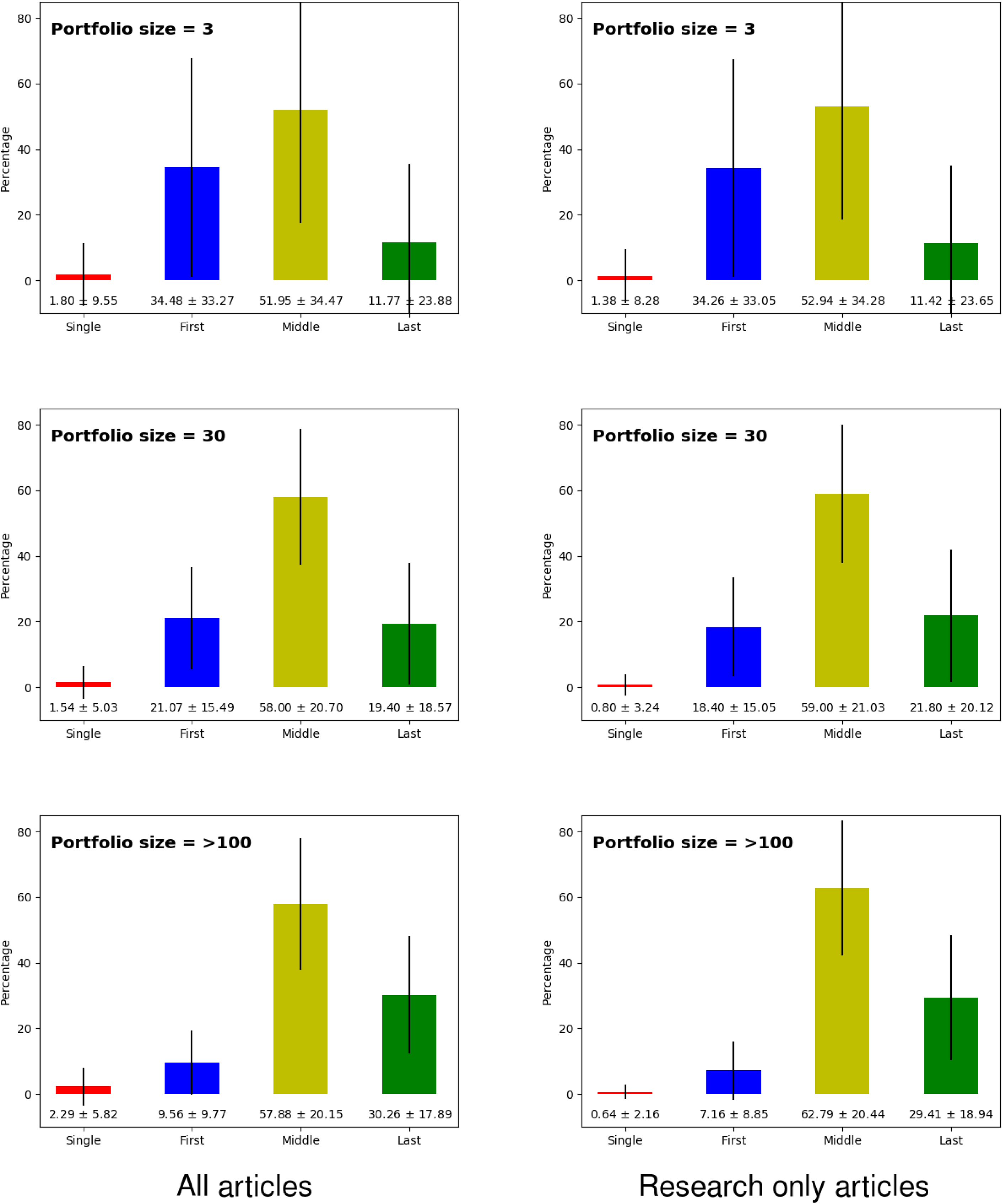
Percentage of single, first, middle and last authorship papers with respect to the size of the portfolio. For the numerical data, see Supplementary Tables 8–9. Animated version of the the figure is available at: http://fcite.org/stats.html#bar_plot

## DISCUSSION

Over the past fifty years, bibliometrics have become an inherent part of the assessment of scientific progress. Such measures, with all of their pros and cons, will be used regardless of whether we support this approach. This development began with the impact factors defined by Eugene Garfield in the 1950s and peaked with the creation of the H-index in 2005. Despite their multiple shortcomings, bibliometrics can offer an instantaneous and relatively fair assessment of science impact. As many scholars have noted previously, one cannot focus on a single number because it cannot embrace the complexity of a researcher’s work (see also Goodhart’s adage). Therefore, it is not proper to focus, for instance, only on the total number of citations or the H-index (which is itself highly correlated with the total number of citations (11)). As those two metrics have been frequently used by funding bodies (consciously/openly or not), researchers have optimized their behaviour, which has led to citation cartels (12), salami slicing (13), and continual increases in the number of authors per publication and the self-citation rate (14). A so-called “publish-or-perish” culture has emerged. The purported quality of a work is inherited directly from impact factors of the journal immediately after publication. The number of authors of papers is irrelevant because all of them receive full credit. Even if someone were to state that the first or the last/ corresponding authors are more important, the community has already found easy “fix” by adding multiple first authors and last authors. This may sound pessimistic, but some fields have already adapted to this new reality very well. Therefore, no one is surprised when a high-energy physics paper has a few thousand authors. Similar approaches are emerging in other fields. In medicine, which already has one of the highest number of authors per publication, many believe that the data provider should be listed as an author of all subsequent publications even without making any other contribution (15). Recently, there is a growing trend towards establishing consortia or groups containing multiple labs/consortia. While this has the advantage of making collaborations that can have synergistic effects, it also has disadvantages. Usually, such initiatives are based on multi-million-dollar grants, and as the results appear, the whole group (usually a few hundred authors) is assigned as the authors of almost every paper produced by the consortium. The *fCite* proposed here stymies such malicious behaviour by simply dividing the citations (or RCRs) by the number of authors while taking into account an author’s number and position. Obviously, the FLAE models used here are far from perfect, and they cannot replace an experienced assessor who reads publications and is familiar with all the insights of his/her own field, but they are a good starting point and certainly a better solution than using the total number of citations or H-index.

All of this being said, one should be aware of the multiple limitations of *fCite*. First, as *fCite* is based on PUBMED, it is not appropriate for many fields that are not well represented in that database (for instance, computer science or social sciences). Second, fCite currently does not filter out self-citations (under development). Moreover, the author has this far been unable to tag and fix all consortium/group papers. Frequently, such items’ authorship appears as “John Smith, Jan Kowalski, SOME Group”, where the first two are leaders, and the SOME group consists of a hundred or more people whose names appear in the supplementary material. FLAE models will count such cases as papers with three authors (likely to have hundreds of citations designating only those three authors and elevating the scores for the first two). Finally, the last shortcoming of *fCite* is that it accords all citations the same weight, which is a massive simplification. This aspect of bibliometrics is well studied and can be considered from many angles. First, not all citations are equal, as a citation can be positive or negative (where the subsequent authors disagree with the original hypothesis). Then, even if the citation is positive, it may have different meanings depending on the section of the article where it is made (the introduction, the methods or the discussion). Moreover, some citations are more important because they are cornerstones for subsequent research, while others are simply review mentions used briefly in the introduction. The other aspect related to citations is that frequently the most important citations are missing due to journal restrictions (for instance, the entire method section, pivotal for any research, is placed in the supplementary material, which has its own reference list and is not listed in most databases). Many such cases can be handled by semantic methods (16), but this approach remains in its infancy. Other characteristics that are frequently used (but not implemented in *fCite*) are the importance of the journal from which the citation comes (e.g., SJR indicators developed by SCImago (17)). Nevertheless, *fCite* is the first, large-scale method that takes into account the number of authors and their positions (only one-time analyses of specific journals, fields, or nations have been done in the past (18) (19)). Additionally, *fCite* uses RCR scores that are taken from iCite (based on PUBMED). Note that these scores differ from citations in many aspects. First, RCR is intended to capture field relevance (it is normalized with respect to the field’s citation levels). Next, in contrast to citations, which are only additive metrics, RCRs can decrease over time. This aspect, while it has been criticized by some (20), is a very useful and demanded feature of bibliometric metrics (that is also missing in the H-index), as RCR can decline when the work begins to be outdated. The other feature of RCR that should not be overlooked is that this metric gives more weight to newer articles (for instance, ten citations for ten-year-old and two-year-old articles will result in dramatically different RCR scores). This is in aggrement with well recognised observation that the citation over article age is skewed and the most of the citations the publication receives within at the age of 2-5 years (Supplementary Fig. 6).

A highly illustrative example is an analysis of top researchers in comparison to the scores provided by Google Scholar and other sources. Supplementary Table 10 reports a selection of statistics for some successful scientists (many of whom are listed in the Highly Cited Researchers (HCR) list created by *Clarivate Analytics*). It is clear that the H-index or total citation counts can often be misleading, and more comprehensive analysis using multiple bibliometric metrics can help. For instance, while HCR is most likely filtered against easy-to-spot cases such as Scientists B and D, it still frequently includes cases such as Scientists A and F. On the other hand, due its limitations (having >15 so-called “Highly Cited Papers”), some outstanding scientists are overlooked (e.g., Scientist C). Therefore, *fCite* can be a very useful tool for deep profiling of even very similar portfolios (with respect to the H-index or total citation count) with surprising discriminatory power (e.g., compare Scientists M and N).

In summary, *fCite* (available free of charge at fcite.org) is a bibliometric tool that provides versatile metrics that can take into account the number of authors and their position on the authorship list. Hopefully, it will facilitate unbiased comparisons of researchers’ importance when they are competing for limited funding and, consequently, enhance scientific development.

## ACKNOWLEDGMENTS

The author acknowledge the NCBI, NLM, ORCID and iCite teams. Funding: L.P.K. (Solid Scientometrics). The author declares competing interests. The Solid Scientometrics company owned by L.P.K. had been established to separate and protect the intellectual property of this project. Since November 2018 L.P.K. is employed in the Institute of Informatics, University of Warsaw, where his research is focused on the computational biology (the proteomics). The project and the Solid Scientometrics company had been conceived before L.P.K had been employed in the University of Warsaw.

## AUTHOR CONTRIBUTIONS

L.P.K. conceived the project, acquired and analyzed the data, developed the web service, and prepared the manuscript.

## DATA AND MATERIALS AVAILABILITY

The *fCite* web service (fcite.org) is available free of charge. All the data needed to evaluate the conclusions in the paper are presented in the Supplementary Materials and at the *fCite* web site. The raw data come from NIH (iCite) and ORCID databases and are available as stated in the Supplementary Materials.

## Supplementary Text

### Motivation

The number of the authors steadily increase over last 25 years (Supplementary Table 5). Moreover, when we divide the publications into research items (describing the original works) and non-research (e.g., the reviews, the editorials, etc.) we clearly can see that on average research publication require more authors. Given the fact that currently the publications with dozens or even hundreds of authors are common, it is desirable to modify the bibliometric metrics (e.g., citations, H-index) to seize the number of the authors for individual paper.

### The patterns of authorship versus portfolio size

The ratio between fractional and total metric declines as the size of portfolio increase (Supplementary Fig. 1). Regardless of the main metric used (RCR or citation) the trend is stable and the data show that small portfolios have more first author publications. When the portfolio size increases more last author items appear. Depending the fractional model used, the small portfolios have 20-25% of contribution of the author falling below 18% for bigger portfolios. Here, it is interesting to point that regardless of fractional model (FLAE vs FLAE2 vs FLAE3 vs EC) the portfolios with around 40-60 items score virtually identical (which means that for mature scientist with multiple publications is not important which fractional model will be used). One should also take into account the spread of the scores which is huge for small size portfolios and decreases over the number of items, but it is always very significant and comprise at least roughly 20% (Supplementary Fig. 2).

**Supplementary Data 1.** PUBMED data set containing publications with PMID numbers from 7 million to 19 million (json format). http://www.fcite.org/icite_1218_7M-19M.tar.gz or http://dx.doi.org/10.18150/repod.3945420

**Supplementary Data 2.** PUBMED data set containing publications with PMID numbers from 20 million to 32 million (json format). http://www.fcite.org/icite_1218_20M-32M.tar.gz or http://dx.doi.org/10.18150/repod.2195699

**Supplementary Data 3.** ORCID Public Data File (xml format). https://figshare.com/articles/ORCID_Public_Data_File_2018/7234028

---

**Supplementary Figure 1.**
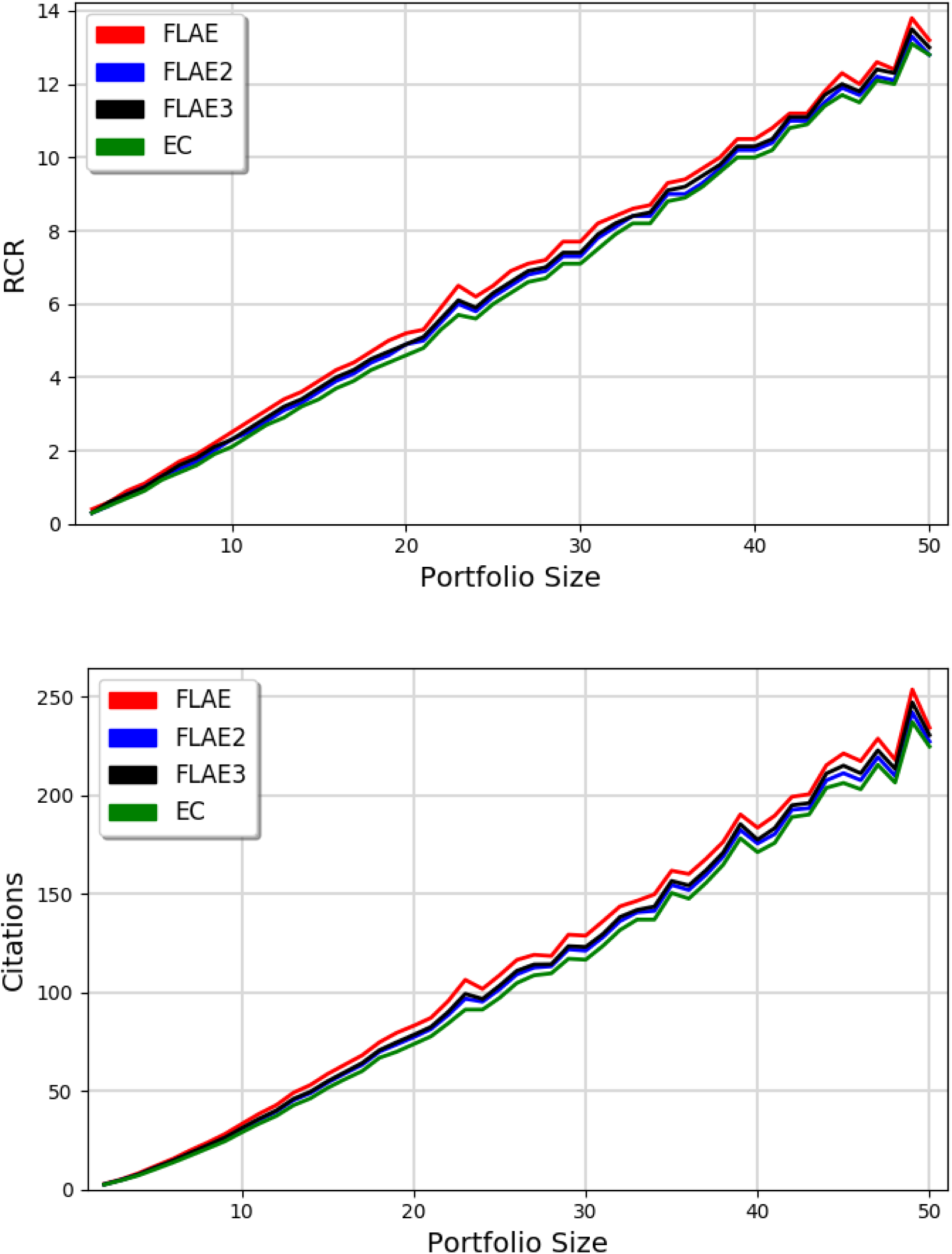
Fractional metrics (FLAE_RCR_, FLAE_Cit_, FLAE2_RCR_, FLAE2_Cit_, FLAE3_RCR_, FLAE3_Cit_, EC_RCR_, EC_Cit_) in respect to the portfolio size (only the 394,189 ORCID portfolios with 2-50 items are presented).

**Supplementary Figure 2.**
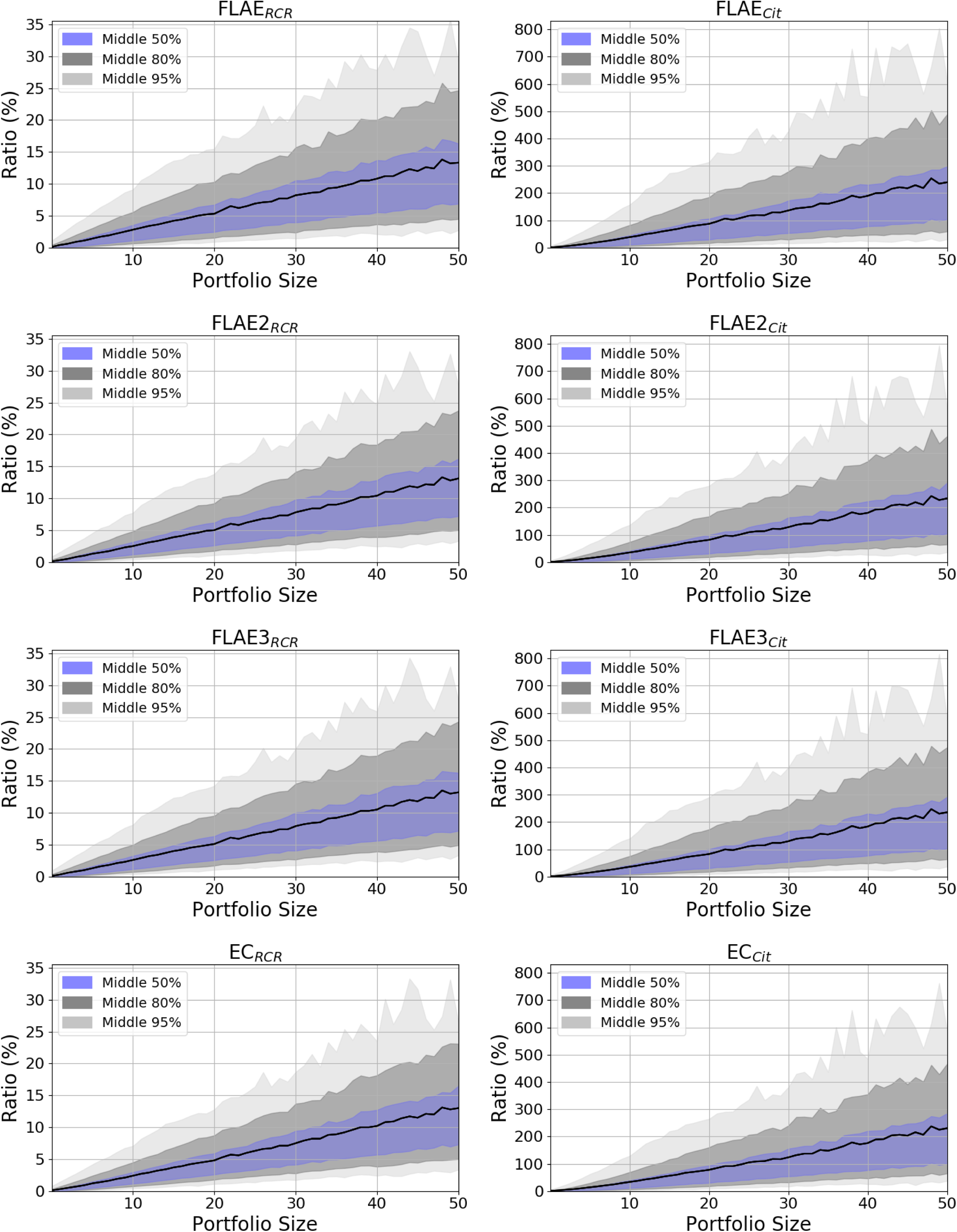
The spread of the fractional metrics (FLAE_RCR_, FLAE_Cit_, FLAE2_RCR_, FLAE2_Cit_, FLAE3_RCR_, FLAE3_Cit_, EC_RCR_, EC_Cit_) in respect to the portfolio size. For the animated plots see fcite.org/stats.html

**Supplementary Figure 3.**
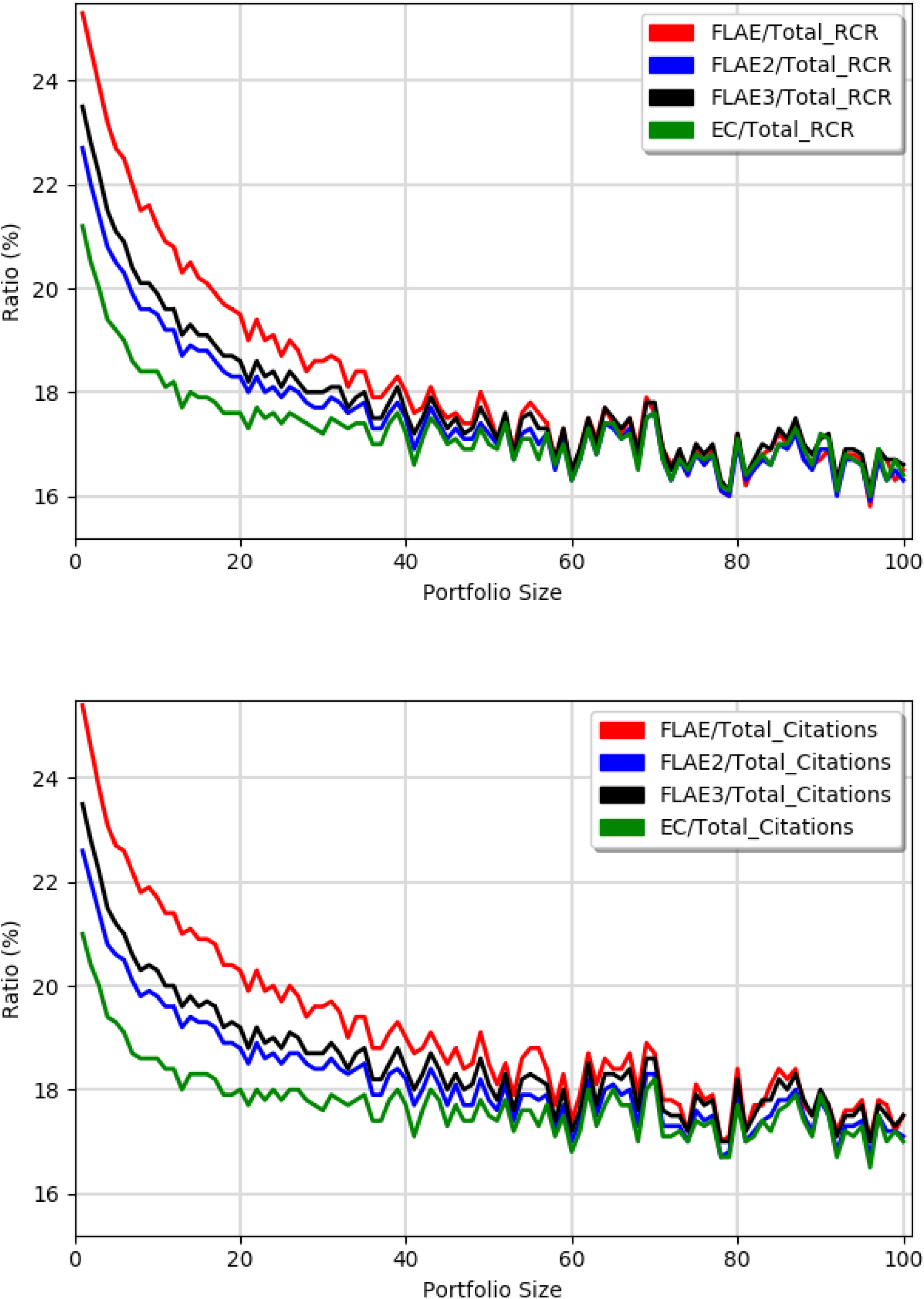
The ratio between the fractional metrics (FLAE, FLAE2, FLAE3, EC) vs. total metrics (RCR or Citations) in respect to the portfolio size.

**Supplementary Figure 4.**
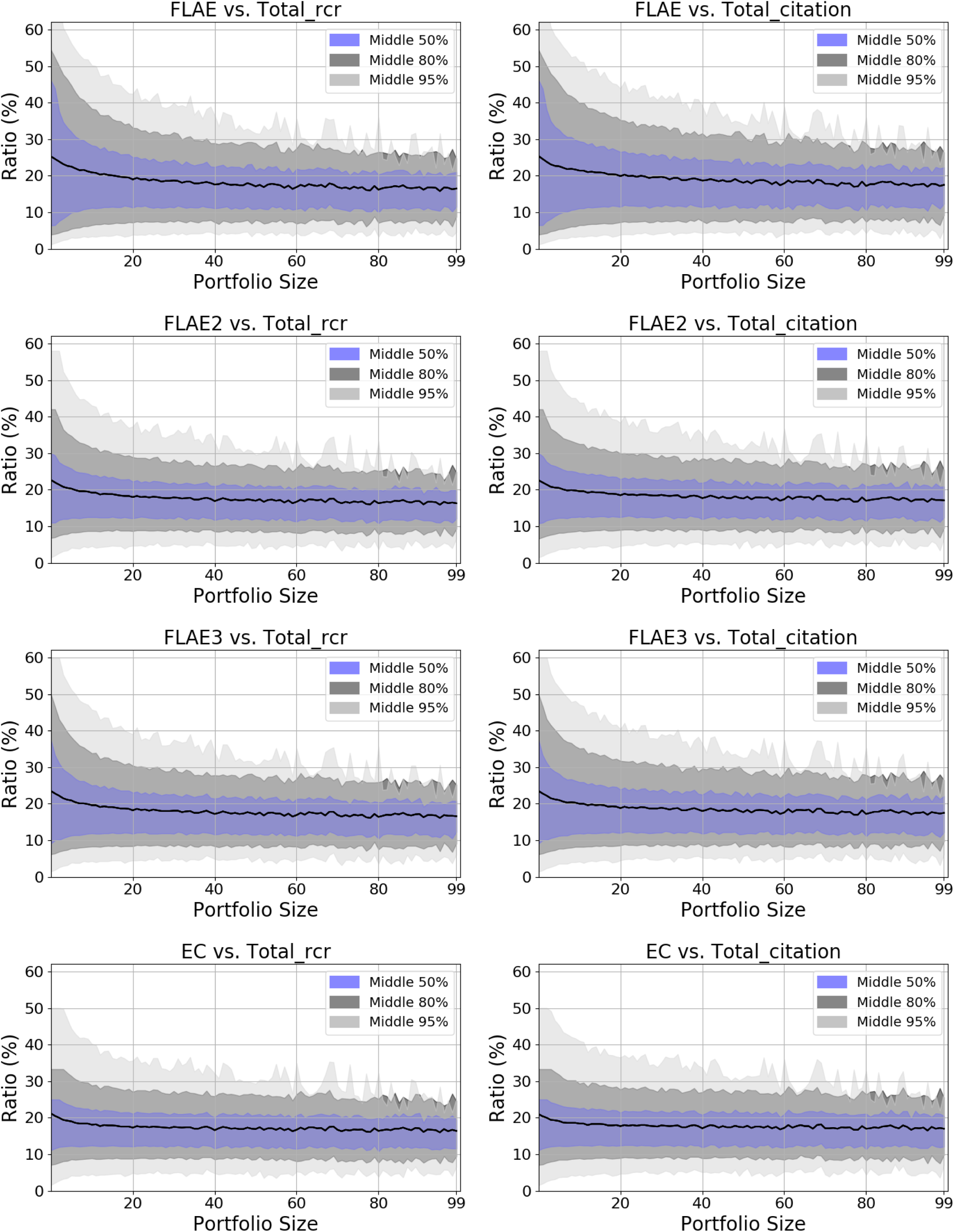
The spread of the ratio between the fractional metrics (FLAE, FLAE2, FLAE3, EC) vs. total metrics (RCR or Citations) in respect to the portfolio size. For the animated plots see http://www.fcite.org/stats.html

**Supplementary Figure 5.**
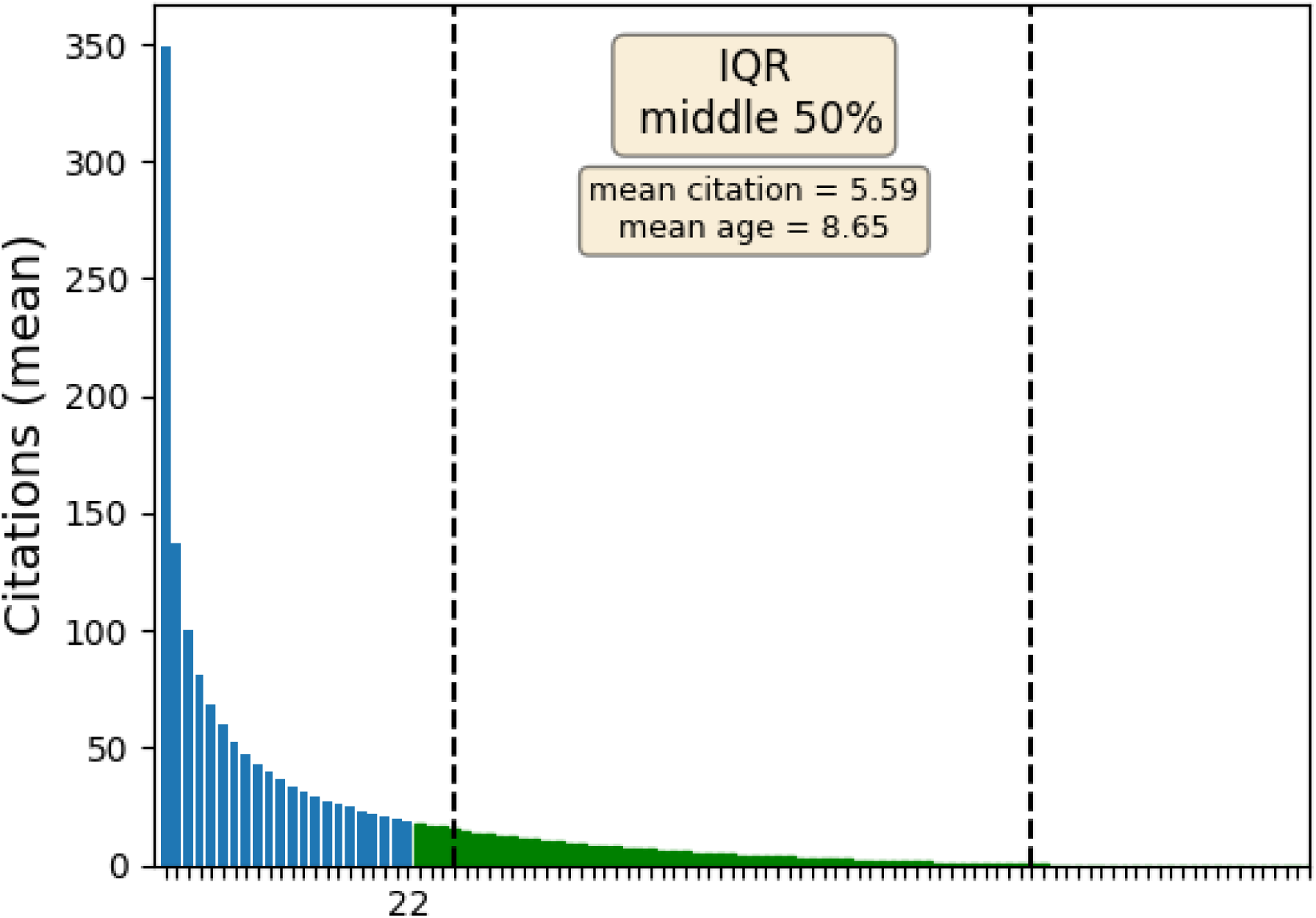
Non-Gaussian distribution of the citations received by the publications (Lotka’s low). Based on 17,787,017 PUBMED publications that received 288,085,850 citations (research and non-research items) within 1995-2018. Blue bars correspond to 80% of the citations (22^nd^ percent of the publications obtained 80% of all citations, in the agreement with Pareto principle). The average publication (within 50% middle range) is expected to obtain only 5-6 citations during 8 years period.

**Supplementary Figure 6.**
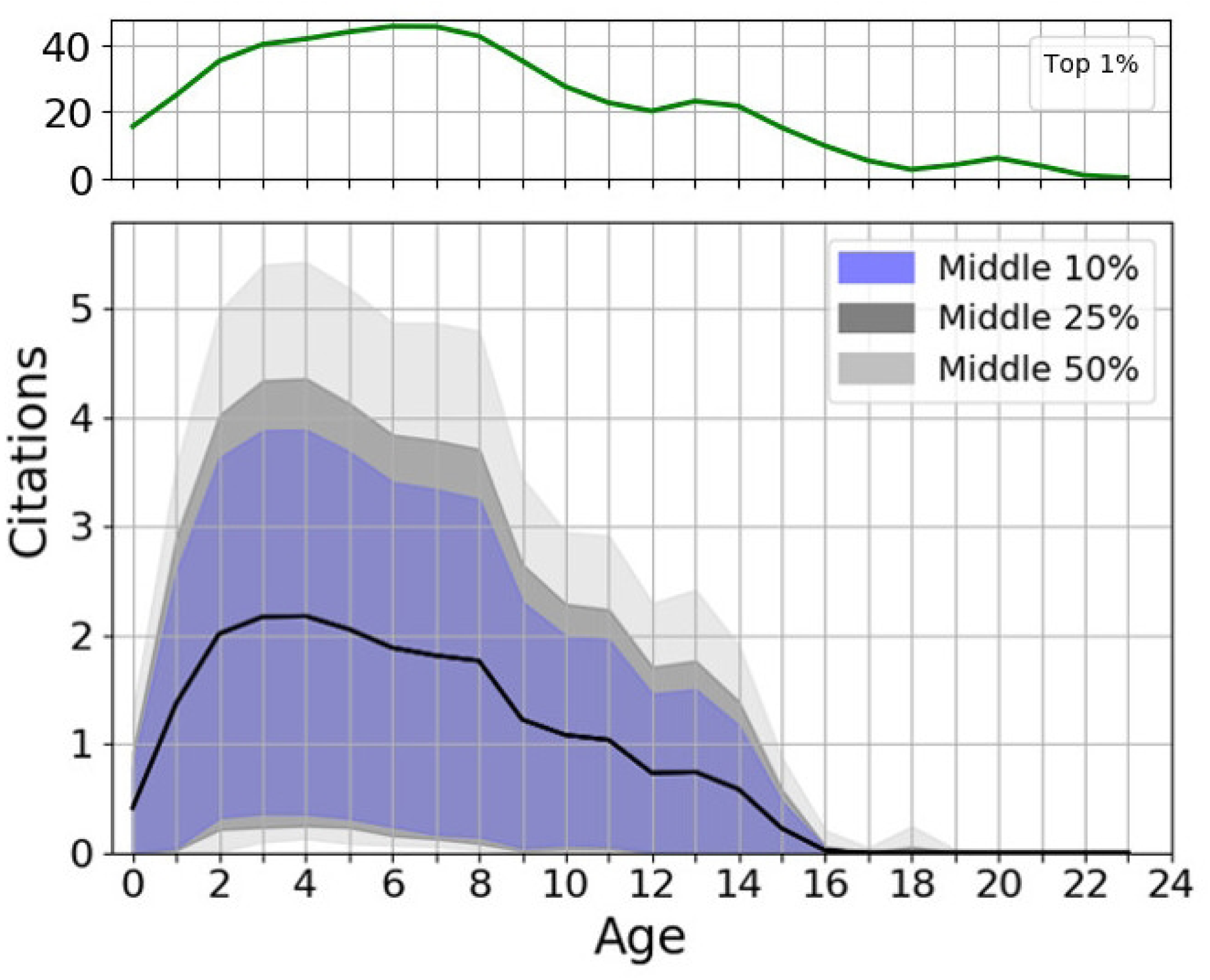
Time dependency of the citations with respect to the age of the article. The citation distribution is highly positively skewed, most of the citations are received within a 2-5 year period followed with long right tail thwarting after 15 years (bottom panel). The similar shape of the distribution can be observed even for the top 1% of best-cited publications, but some differences can be observed. First of all, the number of citations is ~20 times bigger than on average (note different y scales on both panels). Next, the right tail is much longer and the distribution is less skewed.

**SUPPLEMENTARY TABLE 1.** The confusion matrix for up to ten authors for FLAE model. A model with strong emphasis of the first and the last author. For the more details see: http://www.fcite.org/FLAE.txt

| No. Weights for individual authors |  |
| --- | --- |
| 1 | 1.0000 |
| 2 | 0.6667, 0.3333 |
| 3 | 0.5455, 0.1818, 0.2727 |
| 4 | 0.5000, 0.1250, 0.1250, 0.2500 |
| 5 | 0.4762, 0.0952, 0.0952, 0.0952, 0.2381 |
| 6 | 0.4615, 0.0769, 0.0769, 0.0769, 0.0769, 0.2308 |
| 7 | 0.4516, 0.0645, 0.0645, 0.0645, 0.0645, 0.0645, 0.2258 |
| 8 | 0.4444, 0.0556, 0.0556, 0.0556, 0.0556, 0.0556, 0.0556, 0.2222 |
| 9 | 0.4390, 0.0488, 0.0488, 0.0488, 0.0488, 0.0488, 0.0488, 0.0488, 0.2195 |
| 10 | 0.4348, 0.0435, 0.0435, 0.0435, 0.0435, 0.0435, 0.0435, 0.0435, 0.0435, 0.2174 |

**SUPPLEMENTARY TABLE 2.** The confusion matrix for up to ten authors for FLAE2 model (based on (5)). A model with moderate emphasis of the first and the last author. For the more details see: http://www.fcite.org/FLAE2.txt

| No. Weights for individual authors |  |
| --- | --- |
| 1 | 1.0000 |
| 2 | 0.5800, 0.4200 |
| 3 | 0.4200, 0.2700, 0.3100 |
| 4 | 0.3500, 0.2200, 0.1800, 0.2500 |
| 5 | 0.3000, 0.1950, 0.1450, 0.1450, 0.2150 |
| 6 | 0.2600, 0.1700, 0.1300, 0.1200, 0.1200, 0.2000 |
| 7 | 0.2300, 0.1600, 0.1200, 0.1100, 0.1000, 0.1000, 0.1800 |
| 8 | 0.2100, 0.1500, 0.1100, 0.0950, 0.0900, 0.0900, 0.0850, 0.1700 |
| 9 | 0.1950, 0.1400, 0.1000, 0.0850, 0.0820, 0.0810, 0.0790, 0.0780, 0.1600 |
| 10 | 0.1800, 0.1300, 0.0900, 0.0820, 0.0780, 0.0750, 0.0730, 0.0720, 0.0700, 0.150 |

**SUPPLEMENTARY TABLE 3.** The confusion matrix for up to ten authors for FLAE3 model. For the more details see: http://www.fcite.org/FLAE3.txt

| No. Weights for individual authors |  |
| --- | --- |
| 1 | 1.0 |
| 2 | 0.6000, 0.4000 |
| 3 | 0.5000, 0.1667, 0.3333 |
| 4 | 0.4286, 0.1429, 0.1429, 0.2857 |
| 5 | 0.3750, 0.1250, 0.1250, 0.1250, 0.2500 |
| 6 | 0.3333, 0.1111, 0.1111, 0.1111, 0.1111, 0.2222 |
| 7 | 0.3000, 0.1000, 0.1000, 0.1000, 0.1000, 0.1000, 0.2000 |
| 8 | 0.2727, 0.0909, 0.0909, 0.0909, 0.0909, 0.0909, 0.0909, 0.1818 |
| 9 | 0.2500, 0.0833, 0.0833, 0.0833, 0.0833, 0.0833, 0.0833, 0.0833, 0.1667 |
| 10 | 0.2308, 0.0769, 0.0769, 0.0769, 0.0769, 0.0769, 0.0769, 0.0769, 0.0769, 0.1538 |

**SUPPLEMENTARY TABLE 4.**
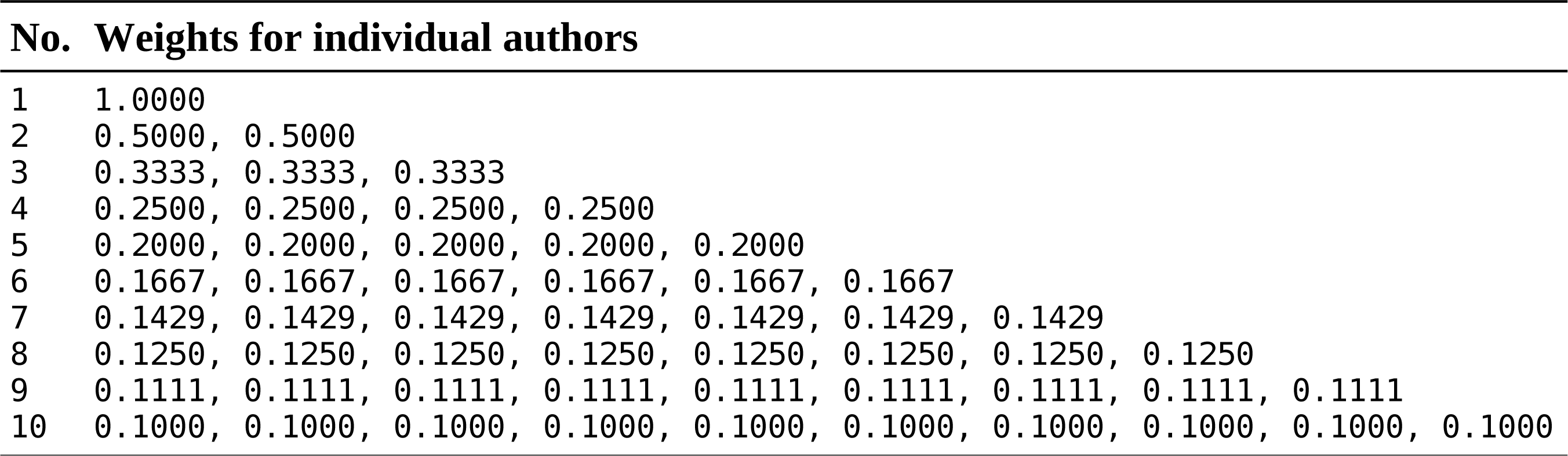
The confusion matrix for up to ten authors for EC model (equal contribution of all authors). For the more details see: http://www.fcite.org/EC.txt

**SUPPLEMENTARY TABLE 5.** The correlations between fractional models and total scores. The lower triangular portion of the matrices (green) correspond to 394,189 ORCID portfolios with 2-50 items and the upper triangular portion of the matrices correspond to all ORCID portfolios with at least single item (600,755 portfolios).

| RCR |  |  |  |  |  | Citations |  |  |  |  |  |
| --- | --- | --- | --- | --- | --- | --- | --- | --- | --- | --- | --- |
|  | FLAE | FLAE2 | FLAE3 | EC | Total |  | FLAE | FLAE2 | FLAE3 | EC | Total |
| FLAE |  | 0.9852 | 0.9929 | 0.9652 | 0.7592 | FLAE |  | 0.9874 | 0.9937 | 0.9708 | 0.7883 |
| FLAE2 | 0.9700 |  | 0.9977 | 0.9951 | 0.7802 | FLAE2 | 0.9741 |  | 0.9980 | 0.9959 | 0.8051 |
| FLAE3 | 0.9884 | 0.9937 |  | 0.9876 | 0.7724 | FLAE3 | 0.9896 | 0.9945 |  | 0.9898 | 0.7985 |
| EC | 0.9237 | 0.9879 | 0.9675 |  | 0.7837 | EC | 0.9363 | 0.9901 | 0.9730 |  | 0.8073 |
| Total | 0.5762 | 0.6166 | 0.6012 | 0.6264 |  | Total | 0.6466 | 0.6850 | 0.6703 | 0.6937 |  |

**SUPPLEMENTARY TABLE 6.** The average number of the authors (PUBMED 17,787,016 publications in 1995-2018).

| Year | Research | Non-research | Research Avg | Not-research Avg | Total Avg |
| --- | --- | --- | --- | --- | --- |
| 1995 | 363007 | 85576 | 3.8849 | 2.0349 | 3.5320 |
| 1996 | 370198 | 87896 | 4.0317 | 2.0976 | 3.6606 |
| 1997 | 363577 | 92127 | 4.1102 | 2.0864 | 3.7011 |
| 1998 | 378481 | 95424 | 4.1766 | 2.0980 | 3.7581 |
| 1999 | 390468 | 102615 | 4.2272 | 2.0834 | 3.7811 |
| 2000 | 421771 | 107840 | 4.2740 | 2.1196 | 3.8353 |
| 2001 | 429734 | 113286 | 4.4378 | 2.1117 | 3.9525 |
| 2002 | 439721 | 118445 | 4.5258 | 2.1878 | 4.0297 |
| 2003 | 457320 | 126100 | 4.6535 | 2.2453 | 4.1330 |
| 2004 | 487038 | 132345 | 4.7827 | 2.3123 | 4.2549 |
| 2005 | 521093 | 134582 | 4.8916 | 2.3505 | 4.3700 |
| 2006 | 545681 | 138391 | 4.9606 | 2.4008 | 4.4427 |
| 2007 | 569964 | 139565 | 5.0582 | 2.4892 | 4.5529 |
| 2008 | 606463 | 143759 | 5.1299 | 2.5105 | 4.6280 |
| 2009 | 638608 | 144718 | 5.2028 | 2.5767 | 4.7176 |
| 2010 | 672589 | 149643 | 5.3194 | 2.6699 | 4.8372 |
| 2011 | 718523 | 156326 | 5.4893 | 2.7427 | 4.9985 |
| 2012 | 775637 | 163479 | 5.6627 | 2.8384 | 5.1710 |
| 2013 | 811789 | 181971 | 5.7698 | 2.9523 | 5.2539 |
| 2014 | 845090 | 195823 | 5.8954 | 3.0620 | 5.3624 |
| 2015 | 878182 | 209943 | 6.0838 | 3.2217 | 5.5316 |
| 2016 | 896432 | 209084 | 6.2214 | 3.4351 | 5.6945 |
| 2017 | 935012 | 189792 | 6.2433 | 3.6815 | 5.8111 |
| 2018 | 927277 | 123246 | 6.1820 | 3.8971 | 5.9139 |
| **Total** | **14444982** | **3342034** | **5.2915** | **2.6970** | **4.8040** |

**SUPPLEMENTARY TABLE 7.** The number of the authors over the years with the respect to the research and non-research items (years 1995-2018).

| Author number | All | Not research | Research | Non research fr | Fr from all |
| --- | --- | --- | --- | --- | --- |
| 1 | 2524210 | 1297946 | 1226264 | 51.4 | 14.2 |
| 2 | 2456770 | 800840 | 1655930 | 32.6 | 13.8 |
| 3 | 2562334 | 477521 | 2084813 | 18.6 | 14.4 |
| 4 | 2429355 | 280806 | 2148549 | 11.6 | 13.7 |
| 5 | 2089498 | 171352 | 1918146 | 8.2 | 11.7 |
| 6 | 1719482 | 111165 | 1608317 | 6.5 | 9.7 |
| 7 | 1203690 | 65535 | 1138155 | 5.4 | 6.8 |
| 8 | 858575 | 41505 | 817070 | 4.8 | 4.8 |
| 9 | 580374 | 25882 | 554492 | 4.5 | 3.3 |
| 10 | 425535 | 18655 | 406880 | 4.4 | 2.4 |
| 11 | 264146 | 11718 | 252428 | 4.4 | 1.5 |
| 12 | 186533 | 8447 | 178086 | 4.5 | 1.0 |
| 13 | 122871 | 6002 | 116869 | 4.9 | 0.7 |
| 14 | 8663 | 4482 | 82211 | 5.2 | 0.5 |
| 15 | 6279 | 3421 | 59373 | 5.4 | 0.4 |
| >15 | 212771 | 16699 | 196072 | 7.8 | 1.2 |

**SUPPLEMENTARY TABLE 8.** Percentage of single-, first-, middle- and last-authorship papers in respect to the size of portfolio for all articles (both research and non-research items). As the number of available portfolios decreases as the size of portfolio increases the data has been bootstrapped 10,000 times (especially crucial for the portfolios with >50 items, where there is less than 1,000 of portfolios passing the threshold).

| Portfolio size | Single | First | Middle | Last | No. portfolios |
| --- | --- | --- | --- | --- | --- |
| 3 | 1.805 ± 9.548 | 34.475 ± 33.269 | 51.954 ± 34.474 | 11.766 ± 23.878 | 45339 |
| 4 | 1.600 ± 8.346 | 33.820 ± 30.054 | 53.406 ± 31.352 | 11.174 ± 21.729 | 33626 |
| 5 | 1.555 ± 7.716 | 33.299 ± 27.893 | 54.422 ± 29.307 | 10.723 ± 20.373 | 26515 |
| 6 | 1.544 ± 7.499 | 32.813 ± 26.068 | 55.360 ± 27.398 | 10.283 ± 19.009 | 21897 |
| 7 | 1.383 ± 6.594 | 32.119 ± 24.739 | 56.157 ± 26.255 | 10.341 ± 18.240 | 18389 |
| 8 | 1.383 ± 6.394 | 31.464 ± 23.650 | 56.643 ± 25.286 | 10.510 ± 17.947 | 16033 |
| 9 | 1.458 ± 6.596 | 31.213 ± 22.931 | 56.310 ± 24.484 | 11.019 ± 17.963 | 13892 |
| 10 | 1.496 ± 6.582 | 30.254 ± 22.022 | 56.988 ± 24.254 | 11.262 ± 17.676 | 12545 |
| 11 | 1.348 ± 5.868 | 29.824 ± 21.263 | 57.296 ± 23.429 | 11.533 ± 17.464 | 10674 |
| 12 | 1.494 ± 6.325 | 29.308 ± 20.851 | 57.367 ± 23.050 | 11.831 ± 17.220 | 9653 |
| 13 | 1.319 ± 5.260 | 28.420 ± 20.173 | 58.037 ± 22.475 | 12.223 ± 17.384 | 8761 |
| 14 | 1.497 ± 5.700 | 28.102 ± 19.678 | 57.624 ± 22.541 | 12.777 ± 17.546 | 7963 |
| 15 | 1.518 ± 5.847 | 27.034 ± 19.400 | 57.899 ± 22.441 | 13.549 ± 17.865 | 7303 |
| 16 | 1.419 ± 5.177 | 26.517 ± 18.750 | 57.879 ± 21.824 | 14.186 ± 18.134 | 6690 |
| 17 | 1.515 ± 6.038 | 26.133 ± 18.512 | 58.186 ± 21.764 | 14.166 ± 17.848 | 6093 |
| 18 | 1.548 ± 5.985 | 25.533 ± 17.812 | 58.381 ± 21.498 | 14.538 ± 17.850 | 5589 |
| 19 | 1.413 ± 5.369 | 25.220 ± 17.959 | 57.909 ± 21.742 | 15.459 ± 18.196 | 5231 |
| 20 | 1.499 ± 5.189 | 24.730 ± 17.712 | 57.797 ± 21.450 | 15.974 ± 18.465 | 4914 |
| 21 | 1.495 ± 5.488 | 23.655 ± 17.131 | 58.381 ± 21.386 | 16.470 ± 18.295 | 4513 |
| 22 | 1.618 ± 5.738 | 23.215 ± 16.769 | 57.678 ± 20.800 | 17.489 ± 18.605 | 4226 |
| 23 | 1.395 ± 4.673 | 22.913 ± 16.182 | 58.659 ± 20.672 | 17.033 ± 18.196 | 3835 |
| 24 | 1.509 ± 5.018 | 22.809 ± 16.454 | 58.410 ± 20.849 | 17.272 ± 18.156 | 3591 |
| 25 | 1.587 ± 5.574 | 21.824 ± 15.937 | 57.868 ± 20.929 | 18.722 ± 19.077 | 3515 |
| 26 | 1.683 ± 5.586 | 21.637 ± 15.999 | 57.910 ± 21.264 | 18.770 ± 18.792 | 3117 |
| 27 | 1.526 ± 4.826 | 21.493 ± 15.934 | 57.873 ± 21.028 | 19.109 ± 19.375 | 3009 |
| 28 | 1.753 ± 5.803 | 20.772 ± 15.170 | 58.115 ± 20.745 | 19.361 ± 18.690 | 2677 |
| 29 | 1.591 ± 5.079 | 21.462 ± 15.548 | 57.553 ± 20.608 | 19.394 ± 18.521 | 2635 |
| 30 | 1.536 ± 5.026 | 21.069 ± 15.493 | 57.997 ± 20.703 | 19.399 ± 18.568 | 2471 |
| 31 | 1.619 ± 5.047 | 20.566 ± 15.645 | 57.736 ± 20.884 | 20.079 ± 18.652 | 2309 |
| 32 | 1.429 ± 4.202 | 19.410 ± 14.861 | 58.121 ± 20.370 | 21.040 ± 18.852 | 2140 |
| 33 | 1.633 ± 5.016 | 19.326 ± 14.633 | 58.249 ± 20.202 | 20.792 ± 18.749 | 2137 |
| 34 | 1.694 ± 4.735 | 19.348 ± 14.385 | 57.530 ± 20.097 | 21.429 ± 18.645 | 1907 |
| 35 | 1.712 ± 5.219 | 19.479 ± 15.002 | 57.346 ± 20.390 | 21.463 ± 18.872 | 1810 |
| 36 | 1.586 ± 4.191 | 18.528 ± 13.804 | 57.546 ± 20.385 | 22.340 ± 18.958 | 1700 |
| 37 | 1.617 ± 4.168 | 18.454 ± 14.539 | 56.617 ± 20.446 | 23.312 ± 19.643 | 1729 |
| 38 | 1.646 ± 4.668 | 17.855 ± 13.562 | 56.644 ± 19.811 | 23.855 ± 19.271 | 1543 |
| 39 | 1.743 ± 4.525 | 17.784 ± 14.104 | 55.639 ± 20.514 | 24.834 ± 19.230 | 1489 |
| 40 | 1.640 ± 5.088 | 18.365 ± 14.100 | 57.916 ± 19.948 | 22.080 ± 18.400 | 1461 |
| 41 | 1.559 ± 4.733 | 17.680 ± 13.569 | 57.068 ± 20.432 | 23.693 ± 18.949 | 1305 |
| 42 | 1.660 ± 4.586 | 17.175 ± 13.710 | 57.428 ± 21.059 | 23.736 ± 19.635 | 1367 |
| 43 | 1.666 ± 3.887 | 17.124 ± 13.465 | 56.138 ± 20.564 | 25.072 ± 19.799 | 1186 |
| 44 | 1.939 ± 6.258 | 16.137 ± 12.661 | 57.008 ± 20.561 | 24.916 ± 19.268 | 1172 |
| 45 | 1.639 ± 3.895 | 16.645 ± 12.841 | 57.957 ± 19.972 | 23.759 ± 18.791 | 1137 |
| 46 | 1.503 ± 3.613 | 16.686 ± 12.620 | 57.234 ± 20.118 | 24.577 ± 19.090 | 1082 |
| 47 | 1.547 ± 4.454 | 16.028 ± 13.354 | 57.871 ± 20.025 | 24.553 ± 19.081 | 1071 |
| 48 | 1.608 ± 4.321 | 16.422 ± 13.348 | 56.813 ± 20.267 | 25.157 ± 19.618 | 995 |
| 49 | 1.961 ± 4.995 | 17.026 ± 14.054 | 54.989 ± 20.002 | 26.023 ± 19.448 | 872 |
| 50 | 1.960 ± 5.016 | 15.880 ± 13.182 | 57.866 ± 21.060 | 24.294 ± 18.893 | 844 |
| 51 | 2.191 ± 6.857 | 15.464 ± 12.934 | 57.460 ± 19.505 | 24.885 ± 18.349 | 833 |
| 52 | 2.015 ± 5.349 | 14.681 ± 12.171 | 57.543 ± 20.084 | 25.760 ± 18.946 | 790 |
| 53 | 1.815 ± 5.731 | 14.966 ± 12.815 | 57.212 ± 19.556 | 26.007 ± 18.938 | 773 |
| 54 | 1.971 ± 5.910 | 15.394 ± 13.198 | 56.118 ± 20.057 | 26.516 ± 19.112 | 714 |
| 55 | 1.963 ± 5.130 | 15.330 ± 12.747 | 55.771 ± 20.303 | 26.936 ± 19.267 | 702 |
| 56 | 1.928 ± 4.848 | 16.255 ± 14.003 | 54.931 ± 20.663 | 26.887 ± 19.520 | 689 |
| 57 | 1.888 ± 4.978 | 14.467 ± 12.267 | 56.794 ± 19.314 | 26.851 ± 18.204 | 665 |
| 58 | 1.715 ± 5.420 | 12.023 ± 11.994 | 63.210 ± 22.986 | 23.053 ± 20.018 | 743 |
| 59 | 2.002 ± 4.978 | 14.772 ± 12.344 | 56.207 ± 20.338 | 27.020 ± 19.037 | 625 |
| 60 | 2.016 ± 5.181 | 13.658 ± 11.762 | 56.921 ± 19.855 | 27.406 ± 19.277 | 596 |
| 61 | 1.997 ± 5.474 | 14.290 ± 12.254 | 56.668 ± 20.230 | 27.046 ± 19.160 | 564 |
| 62 | 2.181 ± 5.761 | 13.878 ± 12.306 | 55.356 ± 20.023 | 28.585 ± 19.948 | 593 |
| 63 | 2.035 ± 5.975 | 13.834 ± 12.589 | 57.733 ± 19.533 | 26.397 ± 18.374 | 513 |
| 64 | 1.942 ± 5.172 | 14.811 ± 12.808 | 55.491 ± 20.259 | 27.756 ± 19.157 | 484 |
| 65 | 2.125 ± 5.615 | 13.566 ± 11.884 | 54.029 ± 20.029 | 30.280 ± 19.507 | 477 |
| 66 | 1.819 ± 3.345 | 13.863 ± 12.170 | 56.334 ± 19.336 | 27.984 ± 18.813 | 480 |
| 67 | 1.987 ± 4.412 | 14.851 ± 11.950 | 54.995 ± 19.528 | 28.168 ± 19.029 | 464 |
| 68 | 1.614 ± 3.076 | 13.586 ± 11.572 | 55.001 ± 19.257 | 29.798 ± 18.689 | 441 |
| 69 | 2.627 ± 7.300 | 13.604 ± 11.599 | 54.233 ± 20.245 | 29.536 ± 19.593 | 443 |
| 70 | 2.094 ± 4.409 | 13.313 ± 10.981 | 54.384 ± 19.327 | 30.209 ± 18.306 | 446 |
| 71 | 2.130 ± 5.856 | 13.517 ± 11.445 | 56.087 ± 20.005 | 28.267 ± 18.747 | 420 |
| 72 | 1.970 ± 4.943 | 13.006 ± 12.200 | 56.742 ± 20.123 | 28.282 ± 18.629 | 378 |
| 73 | 1.964 ± 5.752 | 13.368 ± 12.086 | 56.824 ± 19.408 | 27.845 ± 19.094 | 386 |
| 74 | 1.913 ± 4.216 | 13.206 ± 11.864 | 57.473 ± 19.135 | 27.407 ± 18.432 | 364 |
| 75 | 2.716 ± 9.263 | 12.496 ± 11.310 | 56.160 ± 21.150 | 28.627 ± 18.943 | 389 |
| 76 | 1.756 ± 3.237 | 12.001 ± 9.543 | 56.824 ± 18.841 | 29.420 ± 18.866 | 342 |
| 77 | 1.584 ± 3.346 | 13.373 ± 12.150 | 56.318 ± 19.939 | 28.725 ± 18.627 | 321 |
| 78 | 1.775 ± 4.549 | 10.771 ± 10.555 | 57.105 ± 20.530 | 30.350 ± 19.457 | 332 |
| 79 | 1.963 ± 3.918 | 12.348 ± 10.690 | 59.079 ± 19.993 | 26.609 ± 17.985 | 298 |
| 80 | 2.255 ± 6.448 | 12.759 ± 10.706 | 55.678 ± 19.228 | 29.308 ± 18.376 | 303 |
| 81 | 2.154 ± 4.787 | 11.919 ± 12.474 | 56.828 ± 20.363 | 29.099 ± 18.865 | 289 |
| 82 | 2.307 ± 5.251 | 12.537 ± 10.945 | 55.631 ± 19.251 | 29.525 ± 17.320 | 275 |
| 83 | 2.084 ± 3.601 | 11.856 ± 10.331 | 54.594 ± 19.476 | 31.466 ± 19.081 | 279 |
| 84 | 2.413 ± 5.634 | 13.262 ± 12.046 | 54.214 ± 19.589 | 30.111 ± 17.598 | 250 |
| 85 | 2.906 ± 6.011 | 12.006 ± 11.142 | 53.307 ± 19.023 | 31.781 ± 18.434 | 277 |
| 86 | 1.935 ± 6.884 | 12.048 ± 10.737 | 56.640 ± 18.970 | 29.376 ± 18.088 | 243 |
| 87 | 2.948 ± 8.689 | 12.396 ± 11.480 | 53.846 ± 19.963 | 30.810 ± 18.548 | 267 |
| 88 | 1.876 ± 3.957 | 11.642 ± 11.661 | 55.477 ± 20.183 | 31.005 ± 18.398 | 241 |
| 89 | 2.014 ± 3.353 | 12.898 ± 11.243 | 53.572 ± 18.834 | 31.515 ± 18.705 | 209 |
| 90 | 2.068 ± 5.421 | 11.075 ± 9.559 | 56.342 ± 17.139 | 30.515 ± 17.036 | 204 |
| 91 | 2.726 ± 7.159 | 10.376 ± 9.813 | 54.145 ± 19.719 | 32.753 ± 18.997 | 226 |
| 92 | 1.840 ± 3.979 | 11.475 ± 10.994 | 56.182 ± 19.041 | 30.503 ± 18.615 | 212 |
| 93 | 2.213 ± 3.586 | 12.058 ± 9.917 | 56.018 ± 18.624 | 29.711 ± 18.287 | 177 |
| 94 | 2.574 ± 7.842 | 12.326 ± 10.560 | 57.750 ± 21.297 | 27.351 ± 18.621 | 177 |
| 95 | 2.611 ± 7.487 | 10.869 ± 11.220 | 55.446 ± 20.354 | 31.073 ± 19.047 | 205 |
| 96 | 1.465 ± 2.483 | 10.915 ± 9.385 | 58.132 ± 19.983 | 29.489 ± 19.539 | 207 |
| 97 | 2.873 ± 8.125 | 11.834 ± 10.494 | 56.371 ± 19.763 | 28.923 ± 17.779 | 189 |
| 98 | 1.723 ± 3.703 | 11.758 ± 11.691 | 57.077 ± 19.098 | 29.442 ± 17.300 | 178 |
| 99 | 2.369 ± 5.029 | 10.693 ± 9.744 | 58.345 ± 19.177 | 28.592 ± 17.301 | 157 |
| 100 | 2.252 ± 3.747 | 11.568 ± 10.480 | 55.997 ± 19.712 | 30.183 ± 18.918 | 181 |
| >101 | 2.293 ± 5.823 | 9.565 ± 9.766 | 57.883 ± 20.150 | 30.258 ± 17.890 | 7886 |
|  |  |  |  |  | **Total:** 351849 |

**SUPPLEMENTARY TABLE 9.**
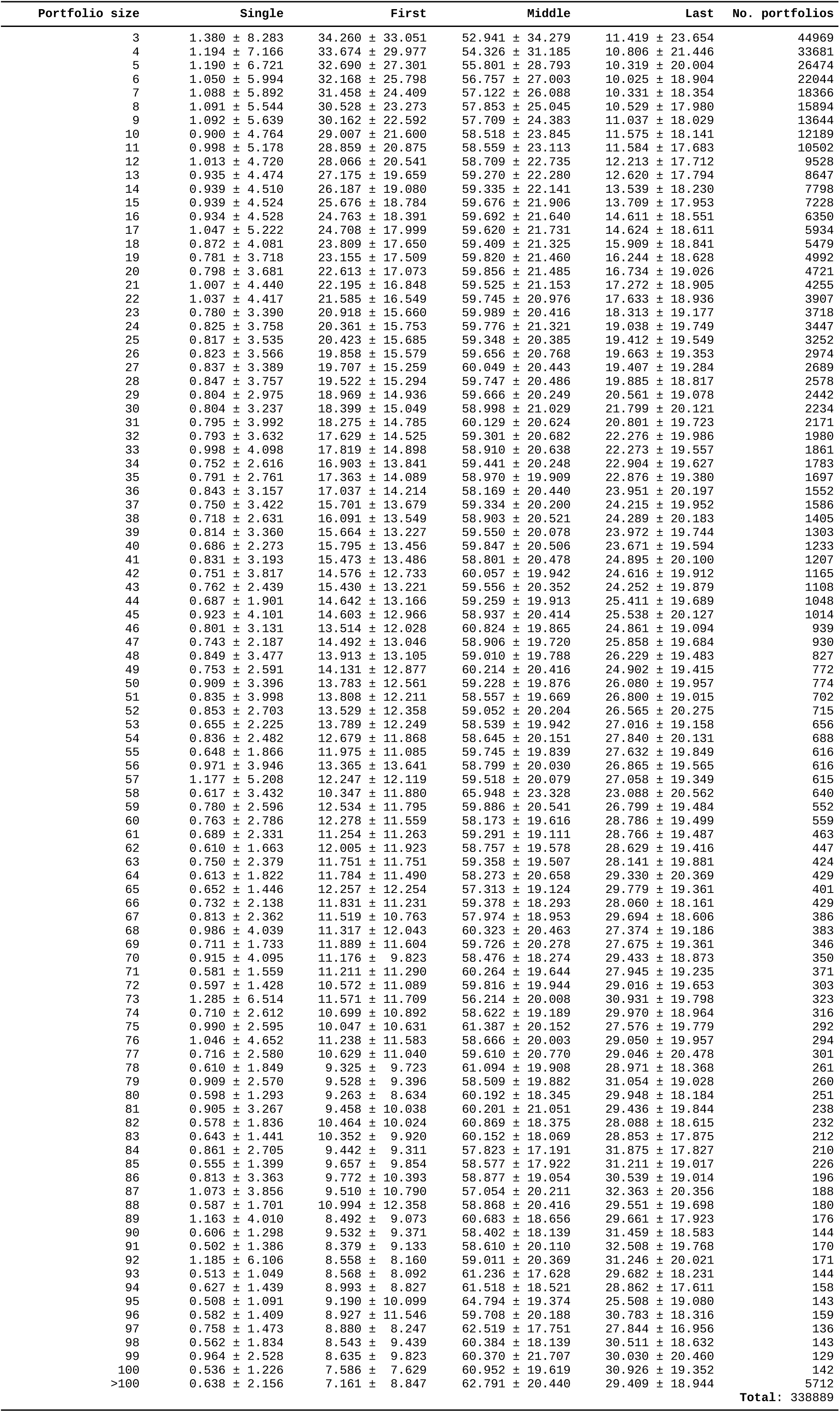
Percentage of single-, first-, middle- and last-authorship papers in respect to the size of portfolio for research articles. As the number of available portfolios decreases as the size of portfolio increases the data has been bootstrapped 10,000 times (especially crucial for the portfolios with >50 items, where there is less than 1,000 of portfolios passing the threshold).

**SUPPLEMENTARY TABLE 10.** A selection of bibliometric statistics for some prominent scientists. For the screenshots from fCite see http://www.fcite.org/examples.html

| Name | Field | In HCR | Articles Number | Avg No Authors | Single % | First % | Middle % | Last % | fCite |  |  |  |  |  | Google Scholar |  |  |
| --- | --- | --- | --- | --- | --- | --- | --- | --- | --- | --- | --- | --- | --- | --- | --- | --- | --- |
|  |  |  |  |  |  |  |  |  | FLAE<sub>RCR</sub> | EC<sub>RCR</sub> | Total RCR | FLAE/RCR | FLAE<sub>Cit</sub> | EC<sub>Cit</sub> | Citations | H-index | Citations |
| Scientist A (ro)<br>Scientist A (all) | Clinical Medicine | 1 | 132<br>163 | 12.7<br>15.8 | 0.00<br>0.61 | 5.30<br>4.29 | 10.61<br>85.28 | 84.09<br>9.82 | 35.5<br>44.2 | 56.7<br>69.7 | 803.4<br>1239.9 | 4.4 %<br>3.6 % | 1017<br>1248 | 1604<br>1952 | 21081<br>32061 | - | - |
| Scientist B (ro)<br>Scientist B (all) | Physics | 0 | 544<br>548 | 1379.3<br>1369.2 | 0.18<br>0.36 | 0.37<br>0.36 | 0.18<br>0.36 | 99.26<br>98.91 | 1.3<br>2.5 | 2.0<br>2.8 | 335.6<br>338.0 | 0.4 %<br>0.7 % | 26<br>54 | 43<br>61 | 2661<br>2717 | 191 | 283,988 |
| Scientist C (ro)<br>Scientist C (all) | Molecular Biology & Genetics | 0 | 33<br>35 | 15.3<br>14.9 | 36.36<br>34.29 | 21.21<br>22.86 | 36.36<br>37.14 | 6.06<br>5.71 | 1087.8<br>1090.1 | 1064.7<br>1025.8 | 1286.7<br>1290.4 | 84.5 %<br>84.5 % | 28429<br>28510 | 27311<br>27372 | 32268<br>32391 | 34 | 60,753 |
| Scientist D (ro)<br>Scientist D (all) | Physics | 0 | 174<br>174 | 2208.9<br>2208.9 | 0.00<br>0.00 | 0.0<br>0.0 | 100.00<br>100.00 | 0.00<br>0.00 | 0.05<br>0.05 | 0.05<br>0.05 | 114.3<br>114.3 | 0.0 %<br>0.0 % | 0.3<br>0.3 | 0.3<br>0.3 | 697<br>697 | 168 | 127,547 |
| Scientist E (ro)<br>Scientist E (all) | Materials Science | 1 | 108<br>112 | 10.2<br>10.0 | 0.00<br>0.00 | 5.56<br>5.36 | 65.74<br>65.18 | 28.70<br>29.46 | 100.0<br>103.9 | 67.8<br>74.8 | 512.7<br>551.7 | 19.3 %<br>18.8 % | 2323<br>2400 | 1497<br>1605 | 11436<br>12037 | 107 | 99,769 |
| Scientist F (ro)<br>Scientist F (all) | Clinical Medicine | 1 | 156<br>296 | 32.4<br>20.6 | 10.26<br>14.19 | 10.90<br>15.54 | 53.85<br>44.93 | 25.00<br>25.34 | 69.5<br>121.1 | 78.6<br>140.2 | 2305.8<br>2769.5 | 3.0 %<br>4.4 % | 1606<br>2402 | 1690<br>2695 | 31927<br>39986 | 102 | 104,919 |
| Scientist G (ro)<br>Scientist G (all) | Physics | 1 | 237<br>246 | 8.1<br>8.0 | 0.42<br>1.22 | 2.38<br>3.25 | 62.03<br>60.57 | 34.18<br>34.96 | 93.4<br>95.8 | 113.9<br>115.7 | 924.9<br>934.7 | 10.1 %<br>10.2 % | 1694<br>1724 | 1820<br>1845 | 14612<br>14735 | 152 | 113,094 |
| Scientist H (ro)<br>Scientist H (all) | Molecular Biology & Genetics | 1 | 185<br>197 | 30.0<br>30.4 | 0.54<br>0.51 | 0.00<br>0.51 | 64.86<br>65.48 | 34.59<br>33.50 | 391.4<br>394.6 | 484.8<br>489.5 | 2949.4<br>2982.0 | 13.3 %<br>13.2 % | 12021<br>12111 | 14808<br>14939 | 90148<br>90864 | 128 | 173,483 |
| Scientist I (ro)<br>Scientist I (all) | Biology & Biochemistry | 1 | 75<br>83 | 19.3<br>18.3 | 0.00<br>1.20 | 14.67<br>16.87 | 85.33<br>80.72 | 0.00<br>1.20 | 106.1<br>110.0 | 84.7<br>87.6 | 939.3<br>948.2 | 11.3 %<br>11.6 % | 2283<br>2372 | 1791<br>1857 | 20596<br>20816 | 71 | 58,008 |
| Scientist J (ro)<br>Scientist J (all) | Biology & Biochemistry | 1 | 296<br>350 | 9.1<br>8.8 | 0.00<br>2.00 | 0.68<br>1.43 | 63.18<br>60.86 | 36.15<br>35.71 | 145.9<br>183.4 | 182.9<br>222.7 | 856.8<br>1022.6 | 17.0 %<br>17.9 % | 4172<br>5520 | 5100<br>6490 | 23419<br>29112 | 115 | 68,249 |
| Scientist K (ro)<br>Scientist K (all) | Chemistry | 1 | 286<br>297 | 7.0<br>6.9 | 0.35<br>0.34 | 0.70<br>1.01 | 59.09<br>59.26 | 39.86<br>39.39 | 111.2<br>131.7 | 132.3<br>149.2 | 748.5<br>848.7 | 14.9 %<br>15.5 % | 2177<br>2523 | 2504<br>2769 | 12871<br>14407 | 128 | 67,952 |
| Scientist L (ro)<br>Scientist L (all) | Chemistry | 1 | 194<br>206 | 8.2<br>8.0 | 0.00<br>0.00 | 2.00<br>1.94 | 55.15<br>55.34 | 42.78<br>42.72 | 65.3<br>73.5 | 56.4<br>67.8 | 394.1<br>448.6 | 16.6 %<br>16.4 % | 1197<br>1284 | 1058<br>1203 | 7313<br>7984 | 87 | 37,474 |
| Scientist M (ro)<br>Scientist M (all) | Immunology | 1 | 187<br>254 | 10.1<br>8.4 | 1.07<br>3.94 | 0.00<br>5.91 | 66.84<br>54.72 | 32.09<br>35.43 | 139.7<br>226.9 | 163.8<br>256.3 | 1011.0<br>1233.2 | 13.8 %<br>18.4 % | 5321<br>9036 | 6451<br>10199 | 34327<br>42998 | 135 | 97,049 |
| Scientist N (ro)<br>Scientist N (all) | Immunology | 1 | 470<br>629 | 17.5<br>14.6 | 0.64<br>1.59 | 1.27<br>3.81 | 74.73<br>64.44 | 23.14<br>30.00 | 79.7<br>153.1 | 68.4<br>140.2 | 1082.3<br>1368.9 | 7.4 %<br>6.9 % | 2492<br>4639 | 1945<br>3892 | 28841<br>36113 | 132 | 65,087 |
| Scientist O (ro)<br>Scientist O (all) | Microbiology | 1 | 243<br>310 | 9.8<br>8.4 | 0.00<br>2.26 | 1.65<br>5.48 | 65.02<br>53.87 | 33.33<br>38.39 | 100.9<br>165.5 | 99.7<br>161.6 | 727.0<br>884.9 | 13.9 %<br>18.7 % | 3085<br>4665 | 3127<br>4558 | 21307<br>24533 | 108 | 44,840 |
| Scientist P (ro)<br>Scientist P (all) | Microbiology | 1 | 485<br>558 | 15.1<br>14.4 | 0.41<br>1.25 | 1.44<br>1.79 | 86.80<br>80.65 | 11.34<br>16.31 | 141.9<br>152.1 | 168.4<br>182.1 | 2488.7<br>2562.0 | 5.7 %<br>5.9 % | 3183<br>3433 | 3318<br>3635 | 51992<br>53533 | 135 | 93,706 |
| Scientist Q (ro)<br>Scientist Q (all) | Plant & Animal Science | 1 | 164<br>191 | 3.8<br>3.6 | 3.05<br>5.76 | 9.15<br>9.95 | 22.56<br>19.90 | 65.24<br>64.40 | 161.6<br>240.8 | 178.2<br>267.1 | 529.0<br>711.1 | 30.6 %<br>33.9 % | 4094<br>6192 | 4502<br>6727 | 13216<br>17729 | 107 | 38,678 |
| Scientist R (ro)<br>Scientist R (all) | Plant & Animal Science | 1 | 135<br>177 | 8.0<br>7.1 | 0.74<br>1.13 | 4.44<br>7.91 | 60.00<br>50.85 | 34.81<br>40.11 | 35.9<br>55.9 | 38.9<br>60.7 | 249.4<br>313.4 | 14.4 %<br>17.8 % | 779<br>1211 | 811<br>1261 | 4771<br>5909 | 66 | 12,896 |

*In HCR* – means that the researcher is included among Highly Cited Researchers 2018 list by Clarivate Analytics *ro/all* – research only or *all* articles

## REFERENCES

1. Garfield, E. (1955) Citation indexes for science; a new dimension in documentation through association of ideas. Science 122, 108–111

2. Hirsch, J. E. (2005) An index to quantify an individual’s scientific research output. Proc. Natl. Acad. Sci. U.S.A. 102, 16569–16572

3. Schmid, S. L. (2017) Five years post-DORA: promoting best practices for research assessment. Molecular biology of the cell 28, 2941–2944

4. Tscharntke, T., Hochberg, M. E., Rand, T. A., Resh, V. H., and Krauss, J. (2007) Author sequence and credit for contributions in multiauthored publications. PLoS Biol. 5, e18

5. Corrêa Jr., E. A., Silva, F. N., da F. Costa, L., and Amancio, D. R. (2017) Patterns of authors contribution in scientific manuscripts. Journal of Informetrics 11, 498–510

6. Lozano, G. A. (2013) The elephant in the room: multi-authorship and the assessment of individual researchers. Current Science 105, 443–445

7. de Solla Price, D. J. (1965) Little science, big science. Columbia University Press New York

8. Jones, M. M., Manville, C., and Chataway, J. (2017) Learning from the UK’s research impact assessment exercise: a case study of a retrospective impact assessment exercise and questions for the future. The Journal of Technology Transfer 1–25

9. Hutchins, B. I., Yuan, X., Anderson, J. M., and Santangelo, G. M. (2016) Relative Citation Ratio (RCR): A New Metric That Uses Citation Rates to Measure Influence at the Article Level. PLoS Biol. 14, e1002541

10. Cohen, W. W., Ravikumar, P., and Fienberg, S. E. (2003) A Comparison of String Distance Metrics for Name-Matching Tasks.

11. Yong, A. (2014) Critique of Hirsch’s citation index: A combinatorial Fermi problem. Notices of the AMS 61, 1040–1050

12. Fister Jr, I., Fister, I., and Perc, M. (2016) Toward the discovery of citation cartels in citation networks. Frontiers in Physics 4, 49

13. Rogers, L. F. (1999) Salami slicing, shotgunning, and the ethics of authorship. AJR. American journal of roentgenology 173, 265–265

14. King, M. M., Bergstrom, C. T., Correll, S. J., Jacquet, J., and West, J. D. (2017) Men set their own cites high: Gender and self-citation across fields and over time. Socius 3, 2378023117738903

15. Longo, D. L. and Drazen, J. M. (2016) Data Sharing. New England Journal of Medicine 374, 276–277

16. Li, D.-C., Liu, H., Chute, C. G., and Jonnalagadda, S. R. (2013) Towards assigning references using semantic, journal and citation relevance.

17. González-Pereira, B., Guerrero-Bote, V. P., and Moya-Anegón, F. (2010) A new approach to the metric of journals’ scientific prestige: The SJR indicator. Journal of informetrics 4, 379–391

18. Abramo, G., D’Angelo, C. A., and Rosati, F. (2013) Measuring institutional research productivity for the life sciences: the importance of accounting for the order of authors in the byline. Scientometrics 97, 779–795

19. Egghe, L. (2008) Mathematical theory of the h-and g-index in case of fractional counting of authorship. Journal of the American Society for Information Science and Technology 59, 1608–1616

20. Janssens, A. C. J., Goodman, M., Powell, K. R., and Gwinn, M. (2017) A critical evaluation of the algorithm behind the Relative Citation Ratio (RCR). PLoS biology 15, e2002536

